# Machine learning enables efficient and effective affinity maturation of nanobodies

**DOI:** 10.64898/2026.01.11.698911

**Authors:** Steffanie Paul, Edward P. Harvey, James Osei-Owusu, Aaron W. Kollasch, Adam J. Riesselman, Conor McMahon, Artem Gazizov, Murali Anuganti, Filmawit Belay, Minh Anh Kieu, Haisun Zhu, L. Robert Hollingsworth, J. Wade Harper, Deborah J. Moshinsky, Andre A. R. Teixeira, Debora S. Marks, Andrew C. Kruse

## Abstract

Antibodies can bind their targets with exquisite potency and selectivity due in part to large antibody-target protein-protein interaction surface areas. Despite the very large size and diversity of synthetic libraries, *in vitro* sorting alone tends to yield binders with modest affinities. By analogy to the *in vivo* affinity maturation in the natural immune system, these initial hits are typically affinity matured *in vitro* to achieve high affinity binding. However, affinity maturation campaigns can be laborious, often requiring multiple selection rounds and strategies for each clone to be optimized. Here, we investigated whether one could accelerate the discovery of optimized binders using machine learning on sequencing data from single selection sorts of affinity maturation yeast-display campaigns. Our results show that sparse sequencing data from a single sorting round can predict sequences that are enriched after multiple rounds. We also find that linear models outperform deep neural networks and semi-supervised approaches in ranking validated affinity-enhancing substitutions. Linear models are also more interpretable, offering insights into residue preferences that can be leveraged for further engineering. We use our models to design and select optimized nanobody binders to relaxin family peptide receptor 1 (RXFP1), yielding multiple improved binders including 3 sub nanomolar binders with the best exhibiting a ∼2500-fold improvement over WT.

## INTRODUCTION

The growing need for effective antibody therapeutics and reagents has led to rapid development in technologies for antibody engineering^1,2^. The antigen-binding properties of an antibody are largely determined by the complementarity determining region (CDR) loops and consequently, these loops are generally the focus of engineering efforts to improve antibody binding^3,4^. A wide variety of antibody formats have been engineered for use as therapeutics and research tools, including single-chain variable fragments (scFvs)^5^, monoclonal antibodies (mAbs)^6^, Fabs, and nanobodies (heavy-chain only camelid antibody fragments)^7,8^. Nanobodies are single domain antibody fragments, and constitute a burgeoning class of clinically relevant biologics with three FDA approved drugs and multiple other candidates in clinical trials^9,10^.

Most FDA-approved antibodies were isolated from immunized animals and then optimized by humanization, improving binding affinity, and improving biophysical properties^2,11^. On the other hand, *in vitro* technologies, such as phage or yeast display, facilitate the identification of binders from synthetically designed libraries. These approaches can provide binding candidates when immunization fails, when there is need to control the composition of the library, and when conformationally selective antibodies are desired^12–18^. *In vitro* technologies and subsequent antibody engineering efforts produced binders with therapeutic properties, as evidenced by the development of the first fully human therapeutic mAb, Humira^19^.

Initial leads from *in vitro* or animal immunization discovery campaigns often need to be optimized in a process called affinity maturation. High-throughput display technologies have been leveraged for affinity maturation, wherein a mutant library of the lead molecule is generated using combinatorial mutagenesis or error-prone PCR (ePCR) and then screened to select for high-affinity variants^1,20^. It is beneficial to generate a large library to maximize chances of finding a high affinity binder; ePCR can produce libraries greater than 10^9^ mutants of a single lead antibody. However, the scale of these campaigns come with added costs. Larger libraries often require many affinity selection rounds to narrow down the pool to a testable number of binders, which requires weeks to months of lab work and significant amounts of lab reagents including large quantities of antigen reagent (>1 mg). Furthermore, successive enrichments increase the chances of experimental artifacts expanding in the library which can confound hit selection. Lastly, hits selected from these campaigns can contain non-affinity-related passenger substitutions that must be pruned out by laborious characterization. Methods that reduce the number of selection rounds required and that identify true optimizing substitutions accurately would greatly improve the efficacy and efficiency of affinity maturation experiments and assist antibody engineers in prioritizing antibodies for follow up studies.

Machine learning (ML) methods that predict the affinity landscape of a specific antibody-antigen interaction have great utility as optimization tools^21–24^. The model predictions are used to select substitutions that would most optimize an initial lead antibody, affording both full control over the diversity of what substitutions are selected and their location in antibodies. Groups have trained ML models on sequencing data from affinity maturation campaigns, using enrichment after sorting as an approximation of binding affinity. These were used to design optimized binders that were shown to have similar or improved affinity in comparison to binders identified experimentally ^21,23^. Notably, both groups trained a model using the enriched sequences from the last FACS round in their campaign to learn the landscape of the highest affinity binders necessitating a full experimental campaign before a model can be trained. In addition, while generative protein models trained on structures have been used to design binders *de novo* without experimental data^25–27^ these are often low affinity and require further optimization^28^. Similarly, although large protein language models (PLMs)^29–31^ have been used to limit the search space of expressible single-substitutions of a starting binding candidate^32^, these models do not consider information about the antigen and thus cannot identify affinity-enhancing substitutions without further laborious validation experiments.

Here, we report a ML framework for designing optimized binders that only requires sequencing data from a single round of FACS affinity sorting. We show that models trained on a single selection round can prospectively predict sequences enriched in further rounds using data from previously completed affinity maturation campaigns. We then used these previously completed affinity maturation campaigns to benchmark model performance, finding that regularized logistic regression models outperform neural network models in identifying single optimizing substitutions; this coupled with the high interpretability of logistic regression models make them powerful tools for optimizing antibodies. Motivated by the ability of the linear regression models to highly rank affinity-enhancing substitutions, we use our framework in a prospective assessment to optimize a nanobody binder to relaxin family receptor I (RXFP1), a clinically relevant GPCR receptor with important roles in regulating cardiac function, angiogenesis, and fibrosis^33^. We use scores from linear models trained on an affinity maturation campaign of this binder to select the best single substitution conferring an over 30-fold increase in affinity. Multiple concurrent amino acid substitutions are oftentimes necessary to achieve desired improvements in antibody potency during affinity maturation campaigns. Furthermore, sometimes antibody variants derived from the same parent that are diverse in sequence space are desired, for instance for broadening patent protections. We also use the models to select sequences with multiple substitutions with respect to the parental clone from the campaign sequencing and design novel sequences using the model in a Gibbs sampling design protocol, yielding 3 subnanomolar binders.

## RESULTS

### Affinity maturation campaigns and data processing

To benchmark our approach, we used next generation sequencing (NGS) data from affinity maturation campaigns for three nanobodies targeting different GPCRs: nanobody AT110, which binds to the human angiotensin II receptor type 1 (AT1R)^34^; nanobody B7, which binds the human β_2_ adrenergic receptor (B2AR); and nanobody RX002, which binds to the relaxin family peptide receptor 1 (RXFP1)^35^. Each nanobody binder was previously discovered via *in vitro* yeast-display selection experiments, starting from a diverse, synthetic nanobody library of approximately 5×10^8^ unique nanobody clones^15^. Nanobody sequences were diversified via ePCR, yielding libraries ranging from 3×10^8^ to 2×10^9^ variants that possessed different numbers of mutations per nanobody. In each case, error-prone PCR libraries were generated and cloned into yeast-display vectors. Each library underwent affinity selection via one MACS round and two or three successive FACS rounds (Supp Figure 1). Nanobody DNA sequences from yeast pools were PCR amplified and sequenced after each selection round. As expected, the number of unique reads decreased with further selection rounds (see Supp Table 1 for complete sequencing statistics). Although we note that many other studies benefit from much deeper sequencing of their libraries^21^, we show here that even with shallower sequencing we are able to extract signal for optimizing substitutions. The predictive ability of sparce sequencing is noteworthy due to practical considerations such as cost savings associated with sparser library sequencing.

Library amplification during NGS sequencing introduces sequencing errors which result in many spurious sequences with very few counts. We reasoned that many of these low-count sequences constituted noise and would confound the ML models. Thus, for each round, we removed any sequence below a count threshold (set to 5 counts) (Supp Table 3). Previous studies have found that sequencing abundance after enrichment correlates poorly with binding affinity^1,36^. One potential explanation for this lack of correlation is bias from sequences that are highly expressed but have modest binding affinity. To account for this, we normalized the abundance after each FACS round by the post-MACS abundance and calculate the resultant log to get a log enrichment ratio (Fig 1B). To normalize the abundance, we would require read counts of the same sequence in both rounds. However, since the overlap of unique reads between sequencing rounds is very low (Supp Table 2) we impute a floor abundance value (set to 1e-6) for any sequence below the count threshold of any round and calculate the enrichment ratio using this value.

**Figure 1:**
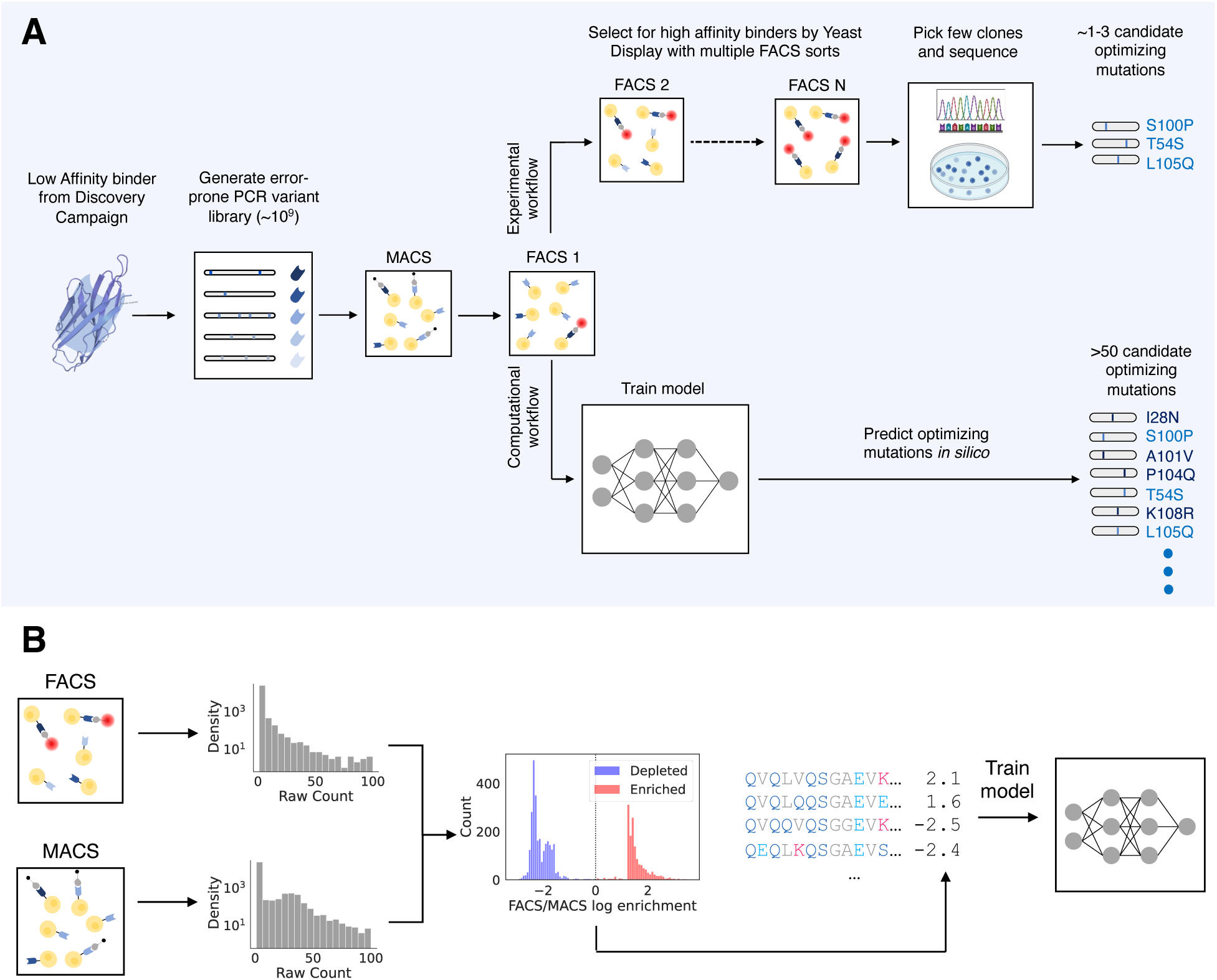
Schematic for our approach. A) Conventional error-prone PCR affinity maturation proceeds with MACS sorting and multiple FACS sorts to filter the library down to a few number of high affinity mutants. Our approach leverages NGS sequencing from a single round of MACS and FACS, coupled with machine learning models to predict optimizing mutations. The model identifies a much larger set of candidate mutations that are selected for downstream validation. B) Data processing pipeline for generating training data. Sequencing counts from the rounds are filtered and normalized to produce an enrichment ratio that acts as a regression target. Inputs to the models are full VHH sequences represented as one hot encodings.

The distributions of log enrichment ratios for the FACS rounds were highly bimodal (Supp Fig 2). This bimodal nature arose because most sequences in the data were only seen in a single sequencing round (Supp Table 2). Thus, in addition to the continuous labels of the log enrichment ratios, we also assigned the binary labels “enriched” and “depleted” to any sequence with log enrichment ratio above and below 0, respectively. We filtered out any sequences for which their sequencing count in both sorts was below the count threshold. Following binary label classification, we aligned each sequence and assigned IMGT numbering to each residue using ANARCI^37^ and only kept IMGT columns that were present in the original nanobody sequence of each campaign. We then one-hot encoded the aligned sequences to produce the inputs to the ML models.

### ML models trained on enrichment from a single FACS round can predict later round enrichment

To see if ML models trained on early selection rounds could predict the enrichment achieved after many selection rounds, we trained models to predict sequence enrichment after FACS1, and tested their predictive ability of FACS2 enrichment. We trained models both for binary classification of the enrichment label and as regression models on the continuous log enrichment value (Fig 2).

**Figure 2:**
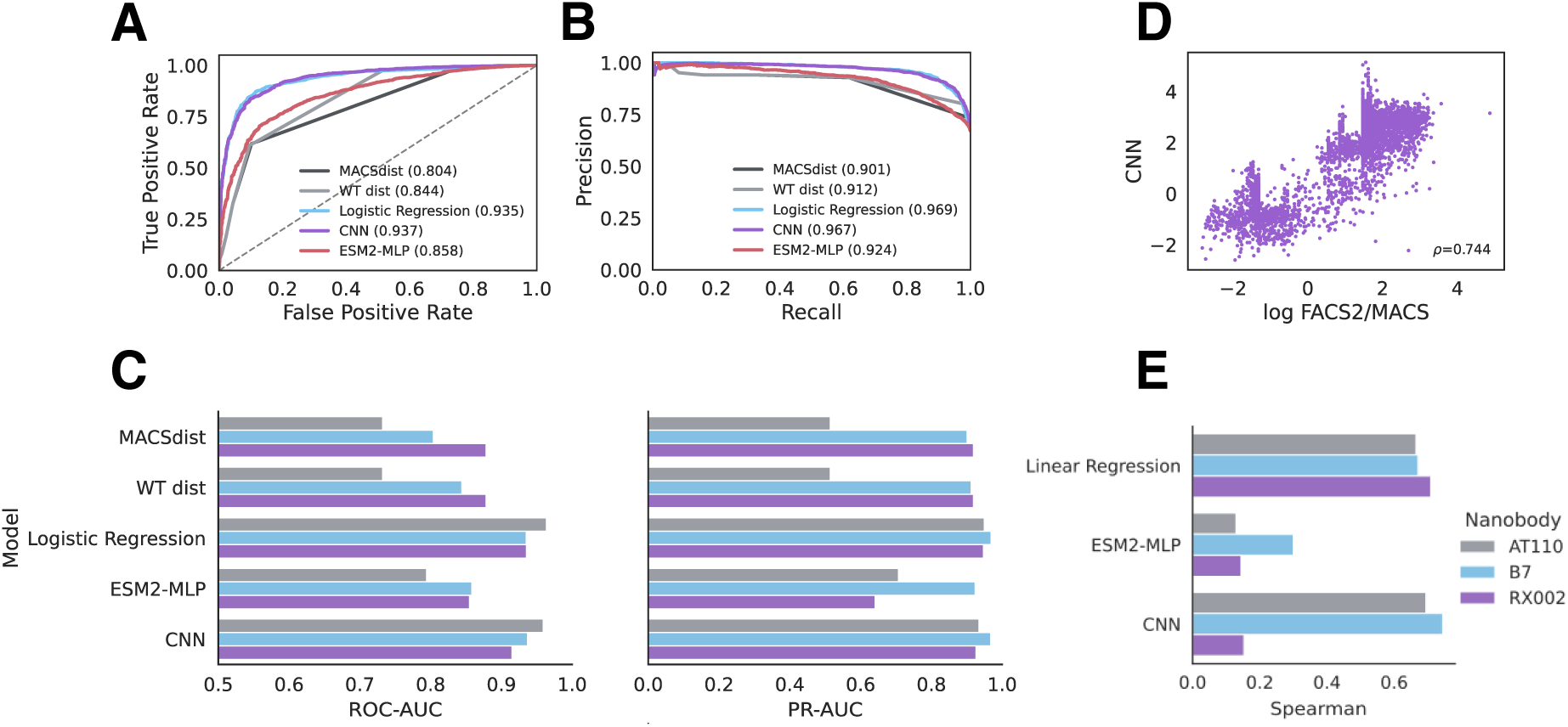
Machine Learning models predict second-round enrichment from first-round data. A) ROC-plots and B) Precision-Recall plots for predicting MACS to FACS2 enrichment by models trained on MACS to FACS1 enrichment from the B7 (β2AR binder) dataset. The area under the curve (AUC) for each model for either metric is shown in brackets in the figure legend. C) Summary performance AT110, B7 and RX002 datasets, evaluated by ROC-AUC (left) and PR-AUC (right). ROC-AUC plots are clipped to 0.5. D) Scatterplot of scores from the top regression model (CNN) on the B7 dataset. E) Spearman correlation of models versus experiment for all datasets.

We trained a variety of models for both regression and binary classification tasks. Linear models with L1 regularization – linear regression and logistic regression for binary classification – have a single weight per amino acid choice for each position in the sequence. Regularization enforces sparsity in the weights such that the non-zero weights in a trained model reflect the most important amino acid substitutions for enrichment/depletion. While linear models are highly interpretable, they cannot model higher-order interactions between residues. Thus, we also trained a convolutional neural network (CNN) which can model nonlinear relationships in the data including residue interactions. Since previous semi-supervised approaches have had some success by leveraging unsupervised learned representations from a large protein language model (PLM) (CITE) we also benchmarked a semi-supervised approach consisting of an MLP top model on top of representations from ESM2, a popular transformer-based PLM ^38^.

The logistic regression and CNN models had similar top performance on both the nanobody B7, AT100 and RX002 datasets. Logistic regression is the highest performing binary classification model on the AT110 data, with an area under the receiver operator curve (ROC-AUC) of 0.964 and an area under the precision recall curve (PR-AUC) of 0.949 (Fig 2 A-C). The same is true of RX002 with an ROC-AUC of 0.935 and a PR-AUC of 0.947. For the nanobody B7 data, logistic regression had slightly lower ROC-AUC and slightly higher PR-AUC than the CNN (Table 1). Comparable performance of the linear logistic regression model with the CNN suggests that higher-order interactions in the sequence may not be required for prediction on this task. However, as our data sizes are small, there may be insufficient data for the neural network models to learn high-order information to improve its prediction. Overall, we find that machine learning models can reliably predict later round enrichment when trained only on a single round of FACS affinity selection after a MACS selection, suggesting there is sufficient affinity information in the sequencing data after a single round to predict enriched binders.

**Table 1:** Summary performance for enrichment and singles prediction. Summary evaluation metrics for the trained machine learning models and sequence distance baselines for the AT110, B7 and RX002 campaigns. ROC-AUC, PR-AUC, NDCG and NDCG@100 were evaluated on classification models. MACSdist refers to the median MACS Hamming distance. Spearman was calculated for regression models. NDCG and NDCG@100 are not reported for sequence distance baselines as all single mutants have the same score under these baselines. NDCG values were not calculated for RX002 as we did not have experimentally validated single substitutions *a priori* to validate the RX002 models.

| **AT110** |  |  |  |  |  |
| --- | --- | --- | --- | --- | --- |
|  | **ROC-AUC** | **PR-AUC** | **Spearman** | **NDCG** | **NDCG@100** |
| *Linear/Logistic Regression* | **0.964** | **0.949** | 0.665 | **0.857** | **0.857** |
| *CNN* | 0.959 | 0.935 | **0.695** | 0.839 | 0.839 |
| *ESM2-MLP* | 0.794 | 0.707 | 0.129 | 0.178 | 0.000 |
| *MACSdist* | 0.732 | 0.515 | - | - | - |
| *WT dist* | 0.732 | 0.515 | - | - | - |
| **B7** |  |  |  |  |  |
|  | **ROC-AUC** | **PR-AUC** | **Spearman** | **NDCG** | **NDCG@100** |
| *Linear/Logistic Regression* | 0.935 | **0.969** | 0.671 | **0.363** | **0.363** |
| *CNN* | **0.937** | 0.967 | **0.744** | 0.255 | 0.235 |
| *ESM2-MLP* | 0.858 | 0.924 | 0.300 | 0.163 | 0.000 |
| *MACSdist* | 0.804 | 0.901 | - | - | - |
| *WT dist* | 0.844 | 0.912 | - | - | - |
| **RX002** |  |  |  |  |  |
|  | **ROC-AUC** | **PR-AUC** | **Spearman** | **NDCG** | **NDCG@100** |
| *Linear/Logistic Regression* | **0.935** | **0.947** | **0.709** | - | - |
| *CNN* | 0.915 | 0.926 | 0.153 | - | - |
| *ESM2-MLP* | 0.855 | 0.641 | 0.144 | - | - |
| *MACSdist* | 0.878 | 0.919 | - | - | - |
| *WT dist* | 0.878 | 0.919 | - | - | - |

As the enrichment labels are highly bimodal and mostly reflect whether the sequence was seen before or after enrichment, we checked that the models were not simply overfitting sequences based on their selection round. To do so, we compared the models to simple distance baselines: i) the Hamming distance from the starting nanobody sequence (WT_dist) and ii) the median Hamming distance to all sequences in the MACS round (median MACS dist). The machine learning models all outperform the distance metric baselines, which suggests that the models are learning some affinity property within the sequences rather than which round the sequences came from.

Of the regression models benchmarked, the CNN has highest Spearman correlation on both the AT110 (0.695) and B7 (0.744) datasets (Fig 2 D-E), marginally outcompeting the linear regression model. However, on the RX002 dataset, the CNN was poorly performant with the linear regression model retaining a higher Spearman of 0.709. While the models show high Spearman correlations across the whole dataset, we found the performance comes from the bimodality of the data; assessing the correlation within just the enriched sequences was much worse (-0.052 for depleted sequences and 0.332 for enriched sequences). In fact, the scores from the CNN regression model have an ROC-AUC and PR-AUC of 0.989 and 0.980, respectively, even higher than the CNN trained as a classification model. This suggests that within an enrichment class, the continuous enrichment values may not separate sequences by affinity with high resolution. This result is expected considering that the FACS gating during affinity selection is stringent and binary, and the abundance of the sequence after or before the gate may not reflect strong differences in binding affinity. This demonstrates the utility of treating prediction of enrichment as a binary classification problem.

Interestingly, for both classification and regression tasks, the semi-supervised approach (ESM2-MLP) had sub-optimal performance, which is logical as the sequences in our datasets are all closely centered around the original nanobody being optimized in the campaign. Thus, evolutionary information from a PLM trained on the protein universe may not be useful for predicting enrichment of these sequences.

Despite the models having high classification performance under ROC-AUC, in comparing scores from a linear regression model against the FACS2/MACS enrichment score, we found that the top binders predicted by the model did not largely overlap with the binders from experimental selections (Supp Fig 7A). This may be because the most enriched binders after FACS2 are simply the highest expressed clones; while we try to account for confounding signal from expression of the sequences by normalizing by the MACS sequencing count, due to the very low overlap between FACS rounds (Supp Table 2) most sequence FACS counts are merely normalized by a constant floor value (1e-6 in our case). The machine learning models generalize information across all the sequences, and thus may be identifying more reliable affinity information, thus leading them to rank other sequences more highly. Interestingly, the sequences ranked by the model have more substitutions in their CDR3 (Supp Fig 7B-C), showing that the models may be ranking more diverse sequences in their top predictions.

### Linear models perform best for recommending single optimizing substitutions

Antibodies are often optimized by identifying single optimizing substitutions that are then combined to produce high affinity binders. While our methods can be used to design binders multiple substitutions away from the initial lead automatically, it may be beneficial to have optimizing information at the single substitutions level as this can easily be combined with domain knowledge and heuristics for avoiding known undesirable properties such as DG or NG isomerization motifs and NxS/T glycosylation sites. Furthermore, of the multiple substitutions found in enriched sequences following a campaign, it is likely that only a small number of these will be true affinity-drivers; we need to know our model can distinguish between substitutions that drive affinity and those that are “passenger” substitutions enriched due to their presence in the same clone as an affinity-enhancing substitution. Thus, we evaluated our methods for how well they can identify single optimizing substitutions.

For initial *in silico* evaluation, we had access to sets of four validated optimizing single substitutions for nanobody B7 and AT110 each. We scored all single substitutions for B7 and AT110 with our models (Fig 3D,E) and evaluated them for how highly they rank the validated substitutions by calculating the normalized discounted cumulative gains (NDCG) of the model’s predictions, weighting each validated substitution equally as a hit. Notably, highly ranked substitutions not in our low-N set may still be real optimizing substitutions; the NDCG does not penalize a model for giving high scores to substitutions not in our set, it merely looks for how highly ranked the validated substitutions are. We also calculated the NDCG when only considering the top 100 ranked substitutions (NDCG@100) as we are often only interested in the top predictions from a model.

**Figure 3:**
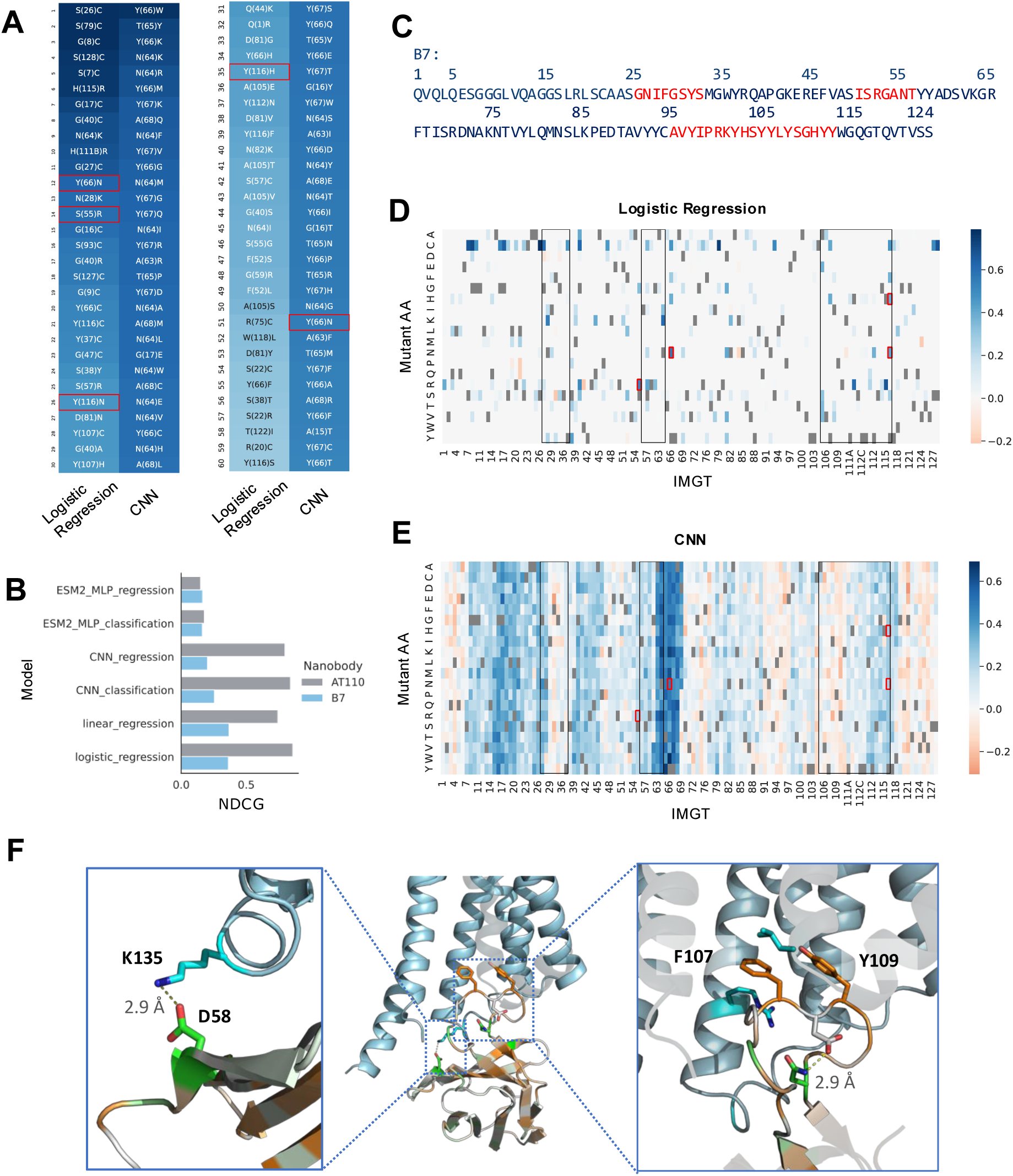
Enrichment prediction models identify optimizing single substitutions. A) The top 60 single substitutions for β2AR binder, B7, ranked by model score for the logistic regression and CNN models. Substitutions are colored by model score (darker blue is higher score) and experimentally validated substitutions are boxed in red. B) NDCG ranking performance for machine learning models for the AT110 and B7 campaigns. C) Amino acid sequence of B7. Indices correspond to residue position in the sequence. CDRs 1-3 are highlighted in red. D-E) Heatmaps showing the gain in score according to the logistic regression model (D) and CNN model (E) for every single substitution of B7. Red boxes correspond to the gain/loss in score for experimentally validated single substitutions and black boxes correspond to CDRs 1-3. (F) (middle) the structure of optimized nanobody AT110i1 bound to the intracellular face of AT1R from PDB:6do1. AT1R is colored in blue and the nanobody is colored by the score of the CNN trained on the AT110 FACS1/MACS enrichment with green corresponding to higher scores (predicted gain in affinity) and orange corresponding to lower scores. (left) a close up of a hydrogen bond formed between D58 in the CDR1 and K135 of AT1R. (right) a close up of the interaction between the CDR3 and the lumen of AT1R. A hydrogen bond within CDR3 residues N113 and D108 is shown. Sticks for F107 and Y109 pointing into the lumen are shown. AT1R residues L404 and R126 are shown as examples of hydrophobic interactions with AT110i1 F107.

We found that logistic regression models had the highest NDCG and NDCG@100 on both the B7 and AT110 validated sets (Fig 3B and Table 1). While the logistic regression and CNN models have similar NDCG on the AT110 data, the logistic regression model identifies all four validated substitutions in the predicted top 15 substitutions, whereas the CNN finds all four in the top 31 substitutions (Supp Fig 4A). This suggests the logistic regression model is meaningfully outperforming the CNN in highly ranking the validated substitutions. As this task only considers single substitutions, it stands to reason that a logistic regression model that does not consider inter-residue interactions would have high performance. Our logistic regression models have a single weight for each single substitution away from the initial lead. L1 regularization during training enforces sparsity on these weights and appropriately accounts for correlation between substitutions, meaning that the model identifies the highest confidence optimizing substitutions for a sequence (Fig 3D). Contrastingly, a CNN model has much less sparsity in its singles prediction (Fig 3E). The model gives high scores for all possible substitutions at certain sites, reflecting that it may be learning which sites may be advantageous to mutate without learning the amino acid preferences necessarily. The corollary of this is that the top singles predicted by the CNN, all cluster around the same sites, leading to low diversity in subsequent selected hits (Supp Fig 4B). In this way, the sparsity in a logistic regression model allows for selecting a highly diverse set of hits across the protein.

### Model scores identify structurally important interacting residues

One of the values of a machine learning model is that it can learn sequence features relating to the specific binding interaction of the nanobody being optimized. To confirm this, we sought to compare the predictions from a model to insights from a structure of a nanobody binding complex. We compared the scores of the CNN trained on the AT110 FACS/MACS enrichment with an available crystal structure for an evolved form of nanobody AT110 (AT110i1) bound to its target, AT1R^34^. We selected the CNN as it had dynamic range in its per-site scores, enabling better qualitative comparison (Fig 3D-E). We colored each residue in the nanobody structure by the max absolute score per site from the model (Fig 3F).

Two of the highest scoring residues from the models were in sites that had evolved from AT110 to AT110i1 (N58D and Y113N – IMGT no. 66 and 115 respectively. D58 in AT110i1 forms a hydrogen bond with site K135 on AT1R (Fig 3F – left), stabilizing the antigen interaction (PDB:6do1)^34^. N113 forms an internal hydrogen bond with D108 (Fig 3F – right) and suggesting that the model learned AT110 substitutions that were important for stabilizing the interaction. The corollary of this, low model scores to mutations of sites 107 and 109 (IMGT no. 112C and 112A respectively), imply that mutating these sites would reduce affinity.

CDR3 residues F107 and Y109 point into the lumen of AT1R and make hydrophobic contacts with multiple lumen residues (including L404 and R126 – shown in Fig 3F (right) – and T61 and A63 – not shown). It is likely that these hydrophobic interactions stabilize the binding complex, supporting the model prediction that a substitution away from these residues would be deleterious. Overall, this structural analysis suggests that our models are learning relevant information relating to the biophysical interaction.

### Interpretable models allow the separation of confounding polyreactivity signal from affinity signal

For nanobody B7 we found that many of the top predictions included cysteines which are known to increase the off-target polyreactivity of an antibody by introducing spurious disulfide bonds (Fig 3D)^39^. This raised concerns that some of the affinity gains from FACS1 and FACS2 may be coming from an enrichment of polyreactive nanobodies. To confirm this, we performed a third FACS sort that included a polyreactivity counter-selection along with the affinity selection (referred to as FACS3 – see Methods). Polyreactivity counter-selection was done using a polyspecificity reagent (PSR) consisting of detergent-solubilized *Spodoptera frugiperda* (Sf9) insect cell membranes as the fluorescence-tagged antigen^40,41^. We trained a logistic regression model on the enrichment between FACS3 and MACS and looked at the weights of the trained model to understand the residue preferences of the FACS3 enrichment. We found that there were fewer cysteine substitutions in the top FACS3 model predictions compared to the models trained on FACS1 and FACS2 enrichment (Supp Fig 5B-C). This confirmed that some of the enrichment of cysteines in FACS1 (Fig3D) and FACS2 (Supp Fig 5A) may have been coming from a population of polyreactive nanobodies in the campaign that were sorted out by the polyreactivity counter-sort. Furthermore, we also found that models trained on the FACS2 enrichment were less predictive of FACS3 enrichment than models trained on FACS1 enrichment by ROC-AUC (Supp Fig 5D). This suggests that there was confounding polyreactivity signal in the selections that was amplified in the library after FACS2. Models trained on this round may be biased towards polyreactivity signal, thus making them less predictive on FACS3 which was experimentally depleted for polyreactive binders. Considering that there may be some polyreactivity bias in our FACS data, when selecting sequences for experimental validation we opted to computationally screen out any substitutions that may increase the binder’s polyreactivity using a pretrained nanobody polyreactivity model developed by our group ^39^. The tool allows us to control the specificity of our binders without requiring further polyreactivity counter-sort experiments.

### Designing and testing novel substitutions for improved nanobody binding affinity

To assess our method prospectively, we predicted the top single substitutions of the RXFP1 ectodomain binder, nanobody RX002, using logistic regression models and validated their affinity experimentally. Activators of RXFP1 may have therapeutic potential for treating cardiovascular disease^42^, and inhibitors may be useful in treatment of reproductive tissue cancers^43^. RX002 was discovered by a yeast-display discovery campaign from a naïve nanobody library against the target RXFP1 ectodomain^15,35^. Logistic regression models were selected as they had the highest singles ranking performance across the models and are more interpretable over CNN models (Fig 3 and Table 1). We scored all possible single substitutions in the CDRs of RX002 with the pretrained polyreactivity model^39^ and considered substitutions that were predicted to not increase the nanobody polyreactivity.

We measured the binding affinity to RXFP1 of the top 11 RX002 substitutions by this selection schema using biolayer interferometry (BLI). 9 out of 11 of the tested substitutions improved the binding affinity of RX002 (Fig 4C, Supp Fig 9, Supp Table 4). Notably, 7 out of 11 of the test substitutions yielded at least 10-fold improvements in binding to the RXFP1 ectodomain. In addition to the negative control nanobody RX002 I51L which was not predicted to improve RX002 affinity, 2 out of the 11 remaining substitutions had a similar K_D_ to RX002. 3 substitutions to residue N27 were in the top ranked predictions (Fig 4E). These 3 substitutions all disrupt an NIS glycosylation site in residues 27 and 28 (Fig 4A). We hypothesized that the glycosylated residue may impede interaction with the target and thus removing it would improve binding. We confirmed that these substitutions eliminated glycosylation by gel electrophoresis and Coomassie blue gel staining. While RX002 showed two bands (for the glycosylated and unglycosylated protein), the mutants only showed one (Supp Fig 8N). Remarkably, all top predicted substitutions were in the CDR1 and CDR3 (except for the negative control substitution I51L in the CDR2 which did not improve affinity). This suggests that either sites in CDR2 do not contribute to target binding and substitutions here do not affect binding; or CDR2 sites are highly important for binding and substitutions disrupt the interaction. Either way, the model predicts that substitutions in CDR2 would not have engineering gains, allowing us to focus attention on CDR1 and CDR3. Overall, these results demonstrate the utility of our methods, and specifically logistic regression models, for identifying single optimizing substitutions.

**Figure 4:**
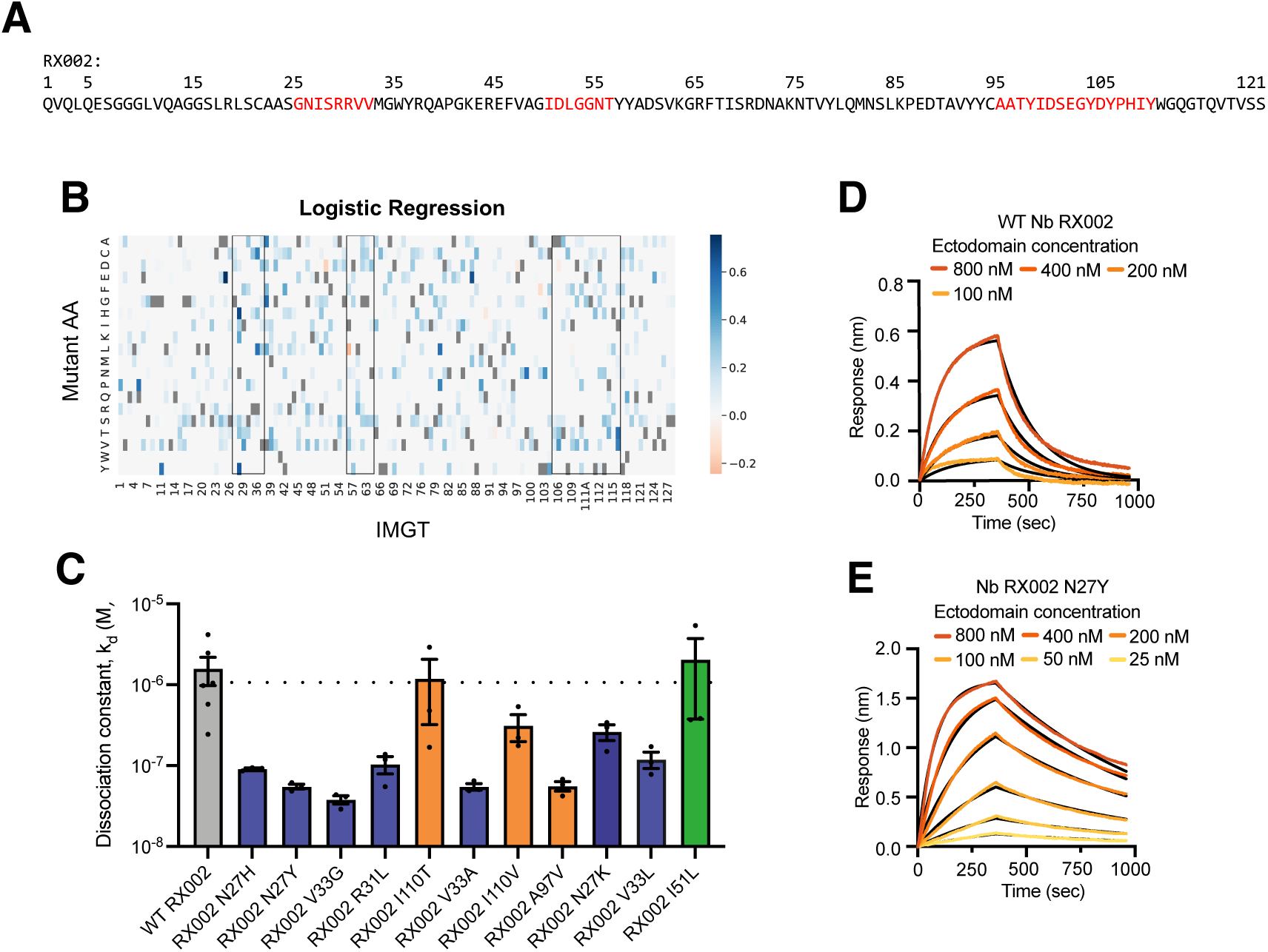
Model selected RX002 mutants have improved binding affinity measured by BLI. A) Sequence of RX002. Red text on the sequence correspond to CDRs 1, 2, and 3. Indices above are according to residue position in the sequence. B) Heatmap of logistic regression scores for all single substitutions for RX002. Black boxes correspond to CDRs 1-3 C) K_D_ for WT RX002 and 10 single substitutions selected using logistic regression model scores. Mutant I51L, a known deleterious mutant^35^ was included as negative control. K_D_ was measured using biolayer inferometry (BLI) at a range of concentrations for 3 biological replicates performed on separate days using separate preparations of antigen and antibody for each substitution. Error bars represent mean +/-SEM for the 3 biological replicates. Dotted horizontal line corresponds to WT k_d_ (average of bioreplicates). Bars for substitutions in CDR1, CDR2 and CDR3 are colored blue, green and orange respectively. C-D) Dissociation curves for WT RX002 (D) and high affinity optimized mutant, N27Y (E) as measured by BLI. Black lines correspond to parametric fit, with the orange gradient corresponding to binder concentration.

### Logistic regression models predict and design sub nanomolar binders

The biggest affinity gains are often achieved by combining multiple optimizing substitutions. A method that can a) select which sequences in the data have the best combination of truly optimizing substitutions and/or b) design novel sequences with ideal combinations has major utility in increasing the success rate of affinity maturation campaigns. We show here that our models are effective in both tasks, enabling major engineering gains.

We used the same logistic regression model trained on the FACS1 and MACS1 selections of the RX002 affinity maturation campaign described above. We used the model to score all the sequences found in the campaign and selected the top 30 sequences with predicted polyreactivity at least as good as WT (LR_score sequences). We also designed novel sequences unseen in the experiment leveraging the logistic regression model in a gibbs sampling procedure (LR_gibbs sequences; see Methods). Of the designs, we selected the top 30 by the model score for validation. Noting that the model tends to prioritize different features that those in the sequences with the highest sequencing counts (Supp Fig 7), we took the top sequences by FACS2 count from the sequencing data, clustered them to reduce sequence redundancy and then selected the top 30 cluster centers by FACS2 count for validation (FACS2_count sequences). These designs were produced as Fc-fusions and the affinity was measured using surface plasmon resonance (SPR; see Methods).

Affinities were successfully measured for 58 candidate designs, with 32 failing to express (Fig 5A). Of those that expressed, the highest affinity binder came from the LR_score set and had a sub nanomolar K_D_ of 0.18 nM. A total of 3 designs had sub nanomolar affinity (two from the LR_score method, one from LR_gibbs and none from the FACS2 sequencing) (Fig 5C,D). Optimizing success can also be considered by the number of designs that are producible and that have affinity greater than that of WT. In this regard, selecting based on FACS2 count had a superior success rate (87%) than the LR_score and LR_gibbs sets (57% and 50% respectively; Fig 5B). Indeed, when considering just the tested binders that were able to be produced, every measured binder from each design method had affinity superior to WT (Fig 5A). From an engineering perspective, the main priority is often finding as high affinity a binder as possible. In this way, the logistic regression model shows excellent performance as a prediction model and a scoring function in a generative process, yielding better sequences than those that could be found from sequencing counts alone.

**Figure 5:**
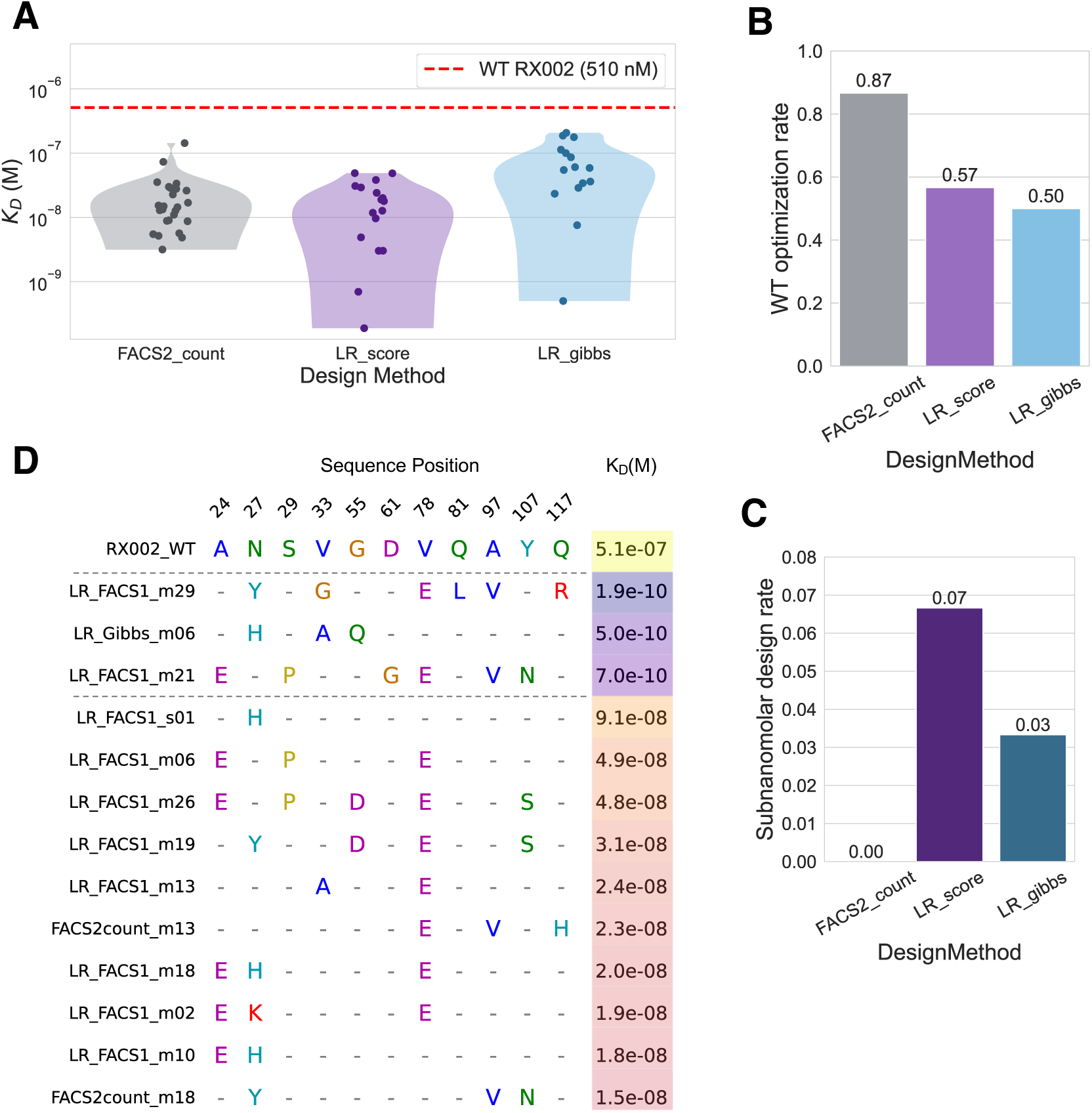
Sequences scored and designed by the logistic regression model reach subnanomolar affinity. A) K_D_s for the tested designs from the 3 methods measured by SPR. Only the K_D_s for successfully measured sequences are shown here (58/90 tests). The affinity of the WT sequence is shown by the red dotted line. B) The fraction of all the tested sequences (including those that did not purify) that had measured affinity greater than the WT sequence by design method. C) The fraction of tested sequences that had affinity in the sub nanomolar range. D) Non-consensus alignment showing a sample of designed positions for the top 13 test sequences (ordered by affinity), with the WT sequence at the top. The amino acids of the WT at the selected positions are shown in full. For tested sequences, the amino acids for any sites containing a substitution with respect to WT are shown, and sites that are the same as WT are just shown as a dash. Dotted black lines demarcate the sub nanomolar designs. The selected sequence positions shown here reflect all the sites in which one of the sub nanomolar designs had a mutation with respect to WT. The K_D_ of each sequence is shown in the right-most column, colored by the log(K_D_) value.

Looking at the sequences with sub-nanomolar affinity, we find that none of the individual substitutions in these sequences fully explains the affinity gains (Fig 5D). Indeed, every individual substitution in these sequences can be found in another sequence with weaker affinity. This suggests that the affinity gains arise from some non-linear relationship between the combination of these substitutions. While the similar performance of the logistic regression model and the CNN in our benchmarking suggested that there was minimal interacting signal between sites in our data, there may still be enough signal to learn ideal combinations in the higher-order mutational landscape explored by the ePCR campaign. This suggests that non-linear models such as CNNs may have even greater design gains. Our workflow combining ePCR libraries of higher-order substitutions with machine learning-enhanced hit-calling, allows us to identify sub-nanomolar affinity conferring combinations of substitutions that a standard ePCR affinity maturation campaign would not. Overall, this establishes the utility of our method as a tool to effectively optimize nanobody affinity.

## DISCUSSION

The growing demand for antibodies as therapeutics and molecular tools has driven the development of faster and cheaper antibody engineering technologies. Affinity maturation is a key part of antibody engineering; however, campaigns are limited by the need for multiple affinity selection sorts to identify a high-quality binder as well as low hit-rates for binder validation after the campaign. We show that machine learning models trained on a single round of FACS sorting from an affinity maturation campaign can replace longer campaigns with multiple FACS sorts for the same property. These models are predictive of i) which sequences would have been enriched in a retrospective analysis of completed campaigns, and ii) which single optimizing substitutions are found after full campaigns. Following experimental binding affinity validation, we found that the models can effectively predict optimizing single substitutions and can design higher-order mutants with vastly increased affinity (Fig 4, Fig 5). Overall, we demonstrate that machine learning models trained on affinity maturation data are useful and efficient antibody engineering tools.

While sequencing counts from post-sort sequencing libraries have not shown to be highly predictive of affinity, machine learning models trained on this data are able to generalize across the sequencing information and learn the salient sequence features that enhance affinity. We find that even with a single round, the model has learned sufficient information to identify substitutions that were only found after a full maturation campaign and multiple validation experiments. This affords antibody engineers the ability to run much shorter affinity maturation campaigns and very quickly identify hits from the campaign using computational methods. Furthermore, by reducing the labor and number of steps required for each selection, our ML framework enables the pursuit of multiple, parallel selection strategies which can yield increased information compared to a single selection. In the B7 campaign, we found that successive rounds were enriching for cysteines which we identified as the enrichment of non-specific binders. This demonstrates a disadvantage in doing longer affinity maturation campaigns with many FACS selections. If there are confounding binders in a library (e.g., highly expressible binders, or very sticky binders), these confounders can be selected for more strongly than on-target, specific binders. Performing many successive sorts risks depleting the library for our on-target binding signal, which would result in many false positives at the end of a campaign. To minimize confounding signal, it may be beneficial to have shorter campaigns which is enabled by our method.

Moreover, this use-case exemplifies the utility of having interpretable models. Being able to interpret a model’s learned information (e.g., by analyzing the weights for each single substitution) allows a practitioner to identify possible confounding signal before hits are validated. Additionally, our modelling framework enables one to flexibly combine domain knowledge (e.g., about residue biases for polyreactivity) with data-driven insights to effectively optimize a binder.

Especially if an engineered antibody is to be used therapeutically, its polyreactivity must be minimized. Affinity and polyreactivity are often correlated, which raises the value in simultaneously optimizing both properties to achieve an optimally specific but high affinity binder. Building computational tools for multi-property optimization is challenging owing to the need for assay data on both properties in the sequence space one wishes to optimize within (e.g., for optimizing an initial lead the models would need to be predictive of the sequence space of possible optimizing mutants around the lead). We and other groups ^36^ enact this by performing joint affinity selection and polyreactivity counterselection experiments on a single library. Towards a more general purpose solution, for cases in which a polyreactivity counter sort was not performed, we leveraged a pretrained polyreactivity model on nanobodies^39^ in combination with our trained affinity models. The polyreactivity predictions did not show high correlation with the affinity models for each campaign (Supp Fig 6) which we reasoned stemmed from the difference in training data for the models. The pretrained polyreactivity model was trained on a much more diverse spread of nanobodies from a naïve library^15^, while our affinity models are trained on sequences from ePCR which are centered around the initial nanobody in sequence space. We opted to use the polyreactivity model as a filter and only selected substitutions that at least retain the initial nanobody specificity for experimental validation. Future work will explore more generalizable models that can be combined across different data sources to construct a single multi-property model to co-optimize antibodies for affinity and specificity.

Computational methods for *in silico* antibody discovery have burgeoned in recent years, with multiple groups reporting methods for discovering VHH^28,44–47^, IgG^44,48,49^, and mini-protein binders ^25,50,51^computationally. While there has been major progress in the field of computational antibody discovery, there is still a way to go before computational methods fully replace the multi-parameter optimization required to convert initial binder hits into potent, selective molecules with drug-like behaviors. Notably, computationally discovered binders had comparable affinities to nanobodies identified by experimental discovery campaigns which are insufficient for most therapeutic uses. Thus, affinity maturation and developability optimization experiments are still necessary to optimize leads (whether identified computationally or experimentally).

Finally, this study focusses on the affinity maturation of nanobodies, but in principle, the methods developed here are fully transferable to any library-level screening of polypeptide binders, e.g., conventional antibodies and non-antibody scaffolds like mini-proteins, DARPins, and short peptides. The fact that the logistic regression models were so powerful also highlights the utility of identifying sparse optimizing substitutions from high-throughput data. Leveraging our modeling framework may also increase the power of screening smaller libraries, as high-value substitutions can be identified before the library is overly bottlenecked. In conclusion, our method is an effective tool to accelerate affinity maturation campaigns of any polypeptide binder.

## STATISTICAL METHODS

Prism software (Graphpad) was used to analyze biolayer interferometry data and perform error calculations. Data are expressed as arithmetic mean ± SEM. Sequencing data analysis and modelling plots generated using the seaborn python package. Models were developed using the scikit-learn and pytorch packages.

## REPORTING SUMMARY

Further information on research design is available in the Nature Portfolio Reporting Summary linked to this article.

## CODE AND DATA AVAILABILITY STATEMENT

The code for training affinity models and predicting affinity enhancing substitutions can be found on github: https://github.com/debbiemarkslab/ML_affinity_maturation.git.

## ACKNOWLEDGEMENTS

This work was funded by a Christopher Walsh Postdoctoral Fellowship to E.P.H.; support by the Howard Hughes Medical Institute through the Hanna H. Gray Fellows program to J.O.-O.; support by the DOE CSGF fellowship DE-FG02-97ER25308 to A.J.R.; NIH grant 5R01NS110395 to J.W.H.; NIH grant 5R01AR079489 to A.C.K.; NIH TR01 grant 5R01CA260415 to D.S.M. and A.C.K; a Chan Zuckerberg Initiative Award (Neurodegeneration Challenge Network, CZI2018-191853) to A.W.K. and D.S.M. We thank Matthew P. Ferguson and Jason G. Liang-Lin for technical assistance, Joseph D. Hurley and Katherine J. Susa for helpful discussions, the Lefkowitz lab and Biswaranjan Pani for providing purified B2AR protein, and the Harvard CMI facility for access to and training on the Octet Red384 BLI instrument.

## AUTHOR CONTRIBUTIONS

S.P., E.P.H., J.O.O., C.M., M.A., F.B., M.A.K., H.Z., L.R.H., D.M., A.A.R.T., A.C.K, and D.S.M. designed experiments, E.P.H., J.O.O., C.M., M.A., M.A.K., F.B., L.R.H. performed experiments; S.P., E.P.H., A.W.K., J.O.O., A.J.R., A.G., M.A., F.B., M.A.K., H.Z., D.M., A.A.R.T., A.C.K., and D.S.M. analyzed data; S.P, AW.K, A.J.R developed the machine learning method; S.P performed all data processing, model training, and computational analysis; M.A., F.B, M.A.K., and H.Z performed SPR measurement of all RX002 multi-substitution variants with supervision from D.M. and A.A.R.T.; A.C.K., D.S.M., A.A.R.T., D.M., and J.W.H. supervised the project and provided funding. S.P., E.P.H., A.C.K., and D.S.M. wrote the manuscript with input from all authors.

## COMPETING INTERESTS STATEMENT

S.P., E.P.H., J.O.O., A.C.K., and D.S.M. are co-inventors on two provisional patent applications for computationally and experimentally obtained anti-RXFP1 nanobodies. S.P. is currently an employee Metaphore Biotechnologies. A.W.K. is currently an employee at Basecamp Research. A.J.R. is currently an employee of Hippo Harvest. C.M. is currently an employee at Sanofi. J.W.H. is a consultant and founder of Caraway Therapeutics (a wholly owned subsidiary of Merck & Co, Inc) and is a member of the scientific advisory board for Lyterian Therapeutics. A.A.R.T. has served and may continue to serve as a consultant for Tectonic therapeutics. D.S.M. is an advisor for Dyno Therapeutics, Octant, Jura Bio, Tectonic Therapeutic, and Genentech, and is a co-founder of Seismic Therapeutic. A.C.K. is a co-founder and consultant for biotechnology companies Tectonic Therapeutic and Seismic Therapeutic, and for the Institute for Protein Innovation, a non-profit research institute. The remaining authors declare no competing interests.

## METHODS

### Generation of error prone PCR libraries

Error prone PCR libraries were constructed to diversify each parent nanobody sequence for affinity maturation (see Supplement for a more detailed protocol). Nucleotide errors were introduced into parent nanobody DNA sequences using the GeneMorph II random mutagenesis kit (Agilent) and the ePCR_MFAlpha_Fwd and ePCR_HA_Rvs primers. To achieve DNA pools with different error rates (ranging from an average of 1 to 6 mutations per nanobody), the amount of template DNA and PCR primer cycles were varied. The error rates of the resulting PCR pools were checked by Gibson cloning nanobody/protein inserts possessing Gibson overlap overhangs added with the ePCR_Lib_fwd and ePCR_Lib_rvs primers into pYDS2.0 vector linearized with the ePCR_Gib_1 and ePCR_Gib_2 primers using NEBuilder HiFi DNA Assembly Master Mix (NEB), transforming the Gibson product into XL1-Blue *E. coli*, and then sanger sequencing 15 random bacterial colonies from the resultant transformations. A second PCR using the ePCR_Lib_fwd and ePCR_Lib_rvs primers was performed on nanobody DNA pools that possessed average error rates greater than 1 to add homology arms needed for yeast homologous recombination. To generate linearized pYDS2.0 vector (which contains a nourseothricin resistance cassette as a selectable marker) for homologous recombination, the pYDS2.0 vector was digested with NheI and BamHI (New England Biolabs), which recognize sites immediately preceding and postceding the nanobody/protein insert sequences, by incubating the vector with the restriction digest enzymes for 1 hour at 37°C. Both the double digested vector and nanobody/protein inserts possessing homology arms were PCR purified and then ethanol precipitated to remove salt. To perform the ethanol precipitation, 0.1x volume 3M sodium acetate pH 5 and 2.5x volume 100% ethanol were incubated with the DNA overnight at -20°C. Following centrifugation at 13,000xRPM for 30 minutes, two sequential washes were performed using 70% ethanol, and DNA pellets were dried at room temperature.

To prepare yeast for library building by homologous recombination, BJ5465 yeast (Addgene) were grown to an OD of 1.5-2, were washed once with water, and then were incubated with 100 mM lithium acetate and 10 mM DTT for 10 minutes at 30°C with shaking. Yeast were washed once more with water and were resuspended in mixture of 5:1 insert to vector in which each 50 mL electroporation reaction contained 5 mg of insert and 1 mg of vector. Electroporations were performed using an ECM 830 Electroporation (BTX-Harvard Apparatus) using square wave pulses at 500V for 15 ms. Following electroporation, yeast were recovered for 45 mins at 30C without shaking in YPAD media. After recovery, dilutions of the transformed yeast were plated on YPAD containing 100 μg/mL nourseothricin to obtain single colony counts to obtain estimates of library diversities. Following dilution plating, 100 ug/mL nourseothricin (Gold Bio) was added to the yeast cultures to select for yeast transformed with nanobody-bearing pYDS2.0 plasmids. Libraries were frozen in aliquots containing at least 10x more yeast cells than the diversity of the library using Yglc4.5 – Trp media supplemented with 10% DMSO.

### Affinity maturation of nanobody binders from error prone PCR libraries

The nanobody B7 β2AR affinity maturation library was expressed in -Trp media containing 2% galactose at 25 °C for two days. Then 2×10^10^ yeast were washed in B2AR selection buffer (20 mM HEPES pH 7.5, 150 mM NaCl, 0.1% LMNG, 0.01% CHS, 10 mM maltose, 0.05% bovine serum albumin) and were pre-cleared using anti-FITC microbeads (Miltenyi) as described above. Following the pre-clear step, yeast were incubated with 500 nM β2AR and 333 nM FITC-anti-FLAG M1 antibody and the LS step was performed as described.

Following the MACS selection, two sequential FACS steps using a SH800 sorter (Sony) were performed at sequentially lower concentrations of β2AR (250 nM and 62.5 nM) to improve the binding affinity of nanobody B7 to β2AR. Then, an affinity selection / polyreactivity counterselection was performed by incubated yeast with PSR reagent (detergent solubilized insect cell membranes) as well as 125 nM FLAG-β2AR and 87.5 nM AF488-anti-FLAG M1 antibody^39^.

Nanobody RX002 was affinity matured to the ectodomain of RXFP1 using a similar workflow (experimental data and methods will be published in a separate manuscript). Briefly, one MACS round was used to deplete the library of nonbinders followed by two sequential FACS rounds at sequentially lower amounts of RXFP1 ectodomain. For each affinity maturation, library selection round aliquots were frozen in Yglc4.5 – Trp media supplemented with 10% DMSO (see Supplement for details).

### Next generation sequencing of nanobody selection rounds

The nanobody sequences were amplified from each selection round via colony PCR. Media was aspirated from 4 x 10^6^ pelleted freshly grown yeast cells per PCR tube in batches of either 16 or 32 samples per selection round. Yeast cells were microwaved dry for 1 min on high power twice, flipping the PCR strip upside down in between microwave rounds. PCR amplification was conducted using 1x Q5 High-Fidelity Master Mix (New England Biolabs) containing 0.3 mM Nanobody_NGS_primer_fwd and Nanobody_NGS_primer_rvs primers containing unique barcodes that allow for selection round deconvolution (see Supplement). PCR amplifications were performed following the manufacturer’s protocol, except for an initial 4 min incubation at 95 °C. Amplified DNA was gel extracted and evaluated via Illumina MiSeq in a 2 x 250 paired-end sequencing reaction (Azenta Life Sciences).

### Recombination protein expression and purification

Parent nanobody RX002, and nanobodies possessing algorithm predicted substitutions introduced by QuikChange Lightning mutagenesis kit (Agilent), were recombinantly expressed as fusions to human IgG1 Fc as described previously^47,52^. First, nanobody DNA sequences followed by a (GGS)3 linker were cloned into pFUSE-CHIg-hG1 (InVivo Gen) containing a H435A substitution in the Fc-domain, which prevents IgG Fc binding to Protein A resin. Since nanobody framework scaffolds also bind Protein A, this construct design ensures quality control since only properly folded nanobody-Fc fusions will bind to the Protein A resin. Then, Expi293F cells (200 mL at a density of 3×10^6^ cells/mL) cultured in Expi293 expression media (Thermo-Fisher) were transiently transfected using nanobody plasmids (0.16 mg) and FectoPro transfection reagent (Polyplus) as a1:1 DNA/FectoPro ratio. After 16 hours, cells were enhanced with 3 mM Valproic acid sodium salt (Sigma-Aldrich) and 0.8% D-(+)-Glucose (Sigma-Aldrich). Transfected cells were boosted with an additional dose of 0.8% D-(+)-Glucose on day 4 and were harvested after 6-7 days once the cell viability decreased below 50%. The media was separated from cells by centrifugation at 4000xg for 15 minutes at 4 °C, filtered using a glass filter, and applied to protein A resin equilibrated with 20 mM HEPES pH 7.5, 150 mM NaCl. The protein A resin was then washed with 20 column volumes 20 mM HEPES pH 7.5, 150 mM NaCl. Following resin washing, nanobody IgG1 Fc fusions were eluted from the resin using 100 mM citrate (pH 3) directly into 2M HEPES pH 8 and the pH of the eluted and neutralized protein solution was checked using pH strips. Eluted nanobody Fc fusions were then dialyzed overnight in 20 mM HEPES pH 7.5, 150 mM NaCl, 10% glycerol and were flash frozen. Protein purity was assessed by SDS-PAGE and analytical size exclusion chromatography runs on a Superdex 200 Increase 3.2/300 gel filtration column (GE Healthcare).

### RXPF1 ectodomain purification

DNA encoding the ectodomain of RXFP1 with an N-terminal hemagglutinin signal sequence and a FLAG tag (DYKDDDDK) was cloned into a pcDNA3.1-Zeo-tetO vector using PCR and NEBuilder HiFi DNA Assembly Mix (New England Biolabs). Site-directed mutagenesis was introduced into plasmids using the QuikChange II XL site-directed mutagenesis kit (Agilent Technologies) and confirmed by Sanger sequencing. FLAG-tagged RXFP1 ectodomain was expressed as a secreted protein in Expi293F cells containing a stably integrated tetracycline repressor (Expi293F tetR, Thermo FisherScientific). Cells were maintained in Expi293 medium (Thermo FisherScientific). The plasmid was transiently transfected into the Expi293F tetR cells using Fectopro. Cells were enhanced 24 hours post-transfection with 0.4% glucose and 3mM sodium valproic acid. Protein expression was induced with 4 μg/mL doxycycline 48 h post-transfection. The supernatant containing the RXFP1 ectodomain was harvested from the cultures 5 days after induction by centrifugation at 4000g for 15 min at 4 °C. To purify the ectodomain, the supernatant was filtered with a glass fiber filter and loaded over M1 anti-FLAG resin equilibrated with 20 mM HEPES pH 7.5 and 300 mM sodium chloride (Buffer A). The resin was washed with buffer A and eluted with buffer A containing 0.2 mg/mL FLAG peptide. The ectodomain was further purified by size exclusion chromatography (SEC) on a Superdex S200 (10/300) Increase column in buffer A. The peak fractions containing RXFP1 ectodomain were collected and concentrated with a centrifugal concentrator with a 10 kDa molecular weight cutoff. Purity of the protein was assessed by SDS-PAGE gel and aliquots were flash frozen in liquid nitrogen and stored at −80°C.

### Biolayer Interferometry experiments

Biolayer Interferometry (BLI) experiments were performed on an Octet RED 384 machine (ForteBio, Menlo Park, CA) at 30 °C. Briefly, Octet AHC2 Biosensors (Sartorius) were pre-wetted with BLI experimental buffer (20 mM HEPES pH 7.5, 150 mM NaCl, 0.02% Tween20) for ten minutes, and then conjugated to Fc-fusion RX002 nanobodies (60 nM) until a loading response of ∼1 nm was achieved. Following a wash for 100 seconds, BLI sensors were then soaked with either BLI experimental buffer or RXFP1 ectodomain at 25 nM, 50 nM, 100 nM, 200 nM, 400 nM, or 800 nM concentrations for 300 seconds to measure associate rate, followed by a 600 second incubation in BLI experimental buffer to measure dissociation rate. K_D_ values were calculated using the BLI Discovery software. To analyze the BLI data, reference sensor (samples with no immobilized ligand) data and reference sample (samples with no analyte) data were subtracted from experimental sensor data. Data were then corrected by aligning the y-axis to the average of the baseline step and performing an interstep correction by aligning data to the dissociation step. Savitsky-Golay noise filtering was then applied to the data. Kinetics analysis was then performed on both the association and dissociation steps using a 1:1 model and global fitting.

### High throughput surface plasmon resonance (HT-SPR)

The SPR binding kinetics experiments were performed on the Carterra LSA platform using HC30M sensor chips (Carterra, cat. #4279) at 25°C. The SPR sensor chip was initially activated by injecting a mixture of 0.033 M N-hydroxysuccinimide (NHS, Millipore Sigma, cat. #130672) and 0.133 M N-(3-dimethylaminopropyl)-N′-ethylcarbodiimide hydrochloride (EDC, Xantec, product code. K AS-50I) in 0.1 M MES buffer (pH 5.5; Carterra, cat. #3625) for 7 minutes. Following activation, goat anti-rabbit IgG Fc was immobilized by injecting a 100 µg/mL solution in 10 mM sodium acetate, pH 5.0, for 10 minutes. Unreacted esters were quenched with 1 M ethanolamine, pH 8.5 (Carterra, cat. #3626) for 10 minutes, resulting in approximately 5000 response units (RU) of immobilization.

Kinetic binding experiments were conducted in a running buffer containing 20 mM HEPES (pH 7.4), 150 mM NaCl, 1 mM CaCl₂, 1 mM MgCl₂, and 0.005% Tween 80. Tested VHH rabbit IgG format antibodies were captured by the pre-immobilized goat anti-rabbit IgG Fc using a 96-channel print-head. Real-time kinetic measurements were acquired by titrating antigen over five concentrations (4.94–400 nM), starting at 400 nM and proceeding via three-fold serial dilutions, with titrations performed from low to high concentrations. For each antigen concentration, data collection times were set to 60 seconds for baseline, 300 seconds for association, and 300 seconds for dissociation. At the end of each antigen titration, the chip surface was regenerated with three 30-second pulses of 0.85% phosphoric acid.

The raw SPR kinetic titration data were analyzed using the Carterra Kinetics software suite with double referencing, which included both reference subtraction (using interspots with no immobilized antibodies) and buffer subtraction (using the leading buffer cycle), followed by data smoothing before kinetic fitting. The kinetic data were fitted to a 1:1 Langmuir binding model, and global fitting parameters for the association rate constant (k_a_, sometimes referred to as the on-rate) and the dissociation rate constant (k_d_, sometimes referred to as the off-rate) were extracted. The equilibrium dissociation constant (K_D_) was calculated as the ratio of the dissociation rate to the association rate (K_D_ = k_d_/k_a_).

### Sequence data processing

#### NGS processing

The paired-end sequencing raw data was QCed and filtered using FastQC and Trimmomatic ^53^. Paired ends were joined using fastq-join. Sequences that do not match the primers used during sequencing were discarded. Nucleotide sequences were translated to amino acid sequences using biopython ^54^. Any protein sequences that did not match (allowing for up to 3 mismatches) the known beginning and end of the initial lead sequence were discarded. Lastly, any sequences with major deletions (>10 AAs) with respect to the initial lead sequence were removed.

#### Enrichment label construction

Low count sequences were reasoned to be the result of sequencing noise and thus sequences with a count less than 5 were removed from each sequencing round. As the abundance of a sequence in any given round may be inflated due to expression biases of each sequence, we normalized the abundance of a sequence in each FACS round by the sequence abundance of the MACS round and took the natural log of this to construct the enrichment value. If a sequence was not found in both rounds (which was the case for most sequences), its log enrichment value was imputed to 1×10^-6^. In addition to the continuous enrichment value, we also binarized the label and set all sequences with enrichment >0 as “enriched” and with enrichment <0 as “depleted”.

#### Sequence preparation

Sequences were aligned using ANARCI ^37^ using IMGT numbering. Only the IMGT columns present in the original lead nanobody were kept as any indels in our sequences probably arose from sequencing errors as our ePCR libraries did not introduce indel variation. The sequences were one-hot-encoded with gaps included (so each site is represented by a 21 length vector). As the elements representing the amino acids for the original lead (WT) are highly correlated in the input, we removed these elements when training logistic/linear regression models to reduce correlation between parameters. In this way each element in the one hot encoding represented a possible substitution from WT.

### Machine learning model training

The code for training affinity models and predicting affinity enhancing substitutions can be found on github: https://github.com/debbiemarkslab/ML_affinity_maturation.git. A variety of machine learning models were benchmarked for this study. Logistic regression, linear regression, CNNs and the top model for the ESM2-MLP semi-supervised model were all trained with L1 regularization the respective loss functions, with weights of 2, 0.0005, 0.0001 and 0.0001 respectively. Neural Network models were trained for 1000 epochs with early stopping. Linear and logistic regression models were built using the implementation in scikit-learn. Neural Network models were built using pytorch. All models were trained on CPU. For the ESM2-MLP model, the ESM2 encodings were retrieved using the pre-trained ESM2_t33_650M_UR50D model using the esm codebase (https://github.com/facebookresearch/esm). The polyreactivity model ^39^ from was implemented using the codebase from the paper (https://github.com/debbiemarkslab/nanobody-polyreactivity).

### Gibbs sampling procedure

To sample novel multiple-mutant sequences unseen in the experiment, scores from the trained ML models were used as energies in a diversity-restricted Gibbs sampling procedure, similar to that used in Fram *et al* 2024^55^. In short, a sampling trajectory is initialized as the WT sequence being optimized (for our experiments this was RX002).

For each step in the trajectory, all single mutants of the current sequence state are scored and combined with a hamming distance regularizing term. The regularized scores are converted to probabilities via a temperature-scaled softmax operation over all single mutants, and the next step is sampled from these probabilities. Noting that the additive nature of the logistic regression models means that more substitutions are always favored, for each sampling trajectory a goal hamming distance from WT is selected, and the energies are penalized to avoid steps that would mutate the sequence further than the goal distance. Substitutions were also restricted to the CDRs of the nanobody to avoid excessive framework substitutions that may disrupt the folding stability of the binder. 100 steps were run for each sample. The sampling trajectories were annealed using a temperature series: (5,1.0,0.5). The temperatures were distributed evenly across the steps.

For a given model and a given WT sequence, 100 sequences were sampled for 3 different goal hamming distances: 3, 5 and 7. The top 30 sequences from each hamming set were selected (yielding 90 designs), and then the top 30 sequences by model score were selected as the final design set.

### NDCG calculation

Normalized discounted cumulative gains (NDCG) is a metric used to measure the quality of a model’s ranking with respect to true scores of the data points (the gains). These scorings can be a true ranking or a binary label. We assigned a gain of 1 to each single substitution in our validated set and a 0 to all others. The NDCG is calculated as:

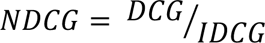

Where the DCG is the discounted cumulative gains (or the score given the rankings from the model) and the IDCG is the ideal DCG (the score given by the ideal ranking in which data points with the highest gains are at the top of the ranking). These are calculated as such:

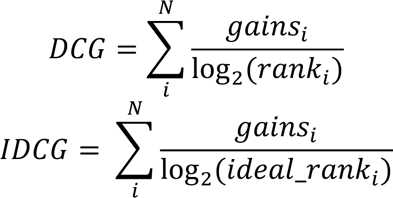

Where the sum is over all single substitutions of the initial lead.

## Notes

https://github.com/debbiemarkslab/ML_affinity_maturation/tree/main

