## Supplementary Figures and Methods for "Machine learning enables efficient and effective affinity maturation of nanobodies"

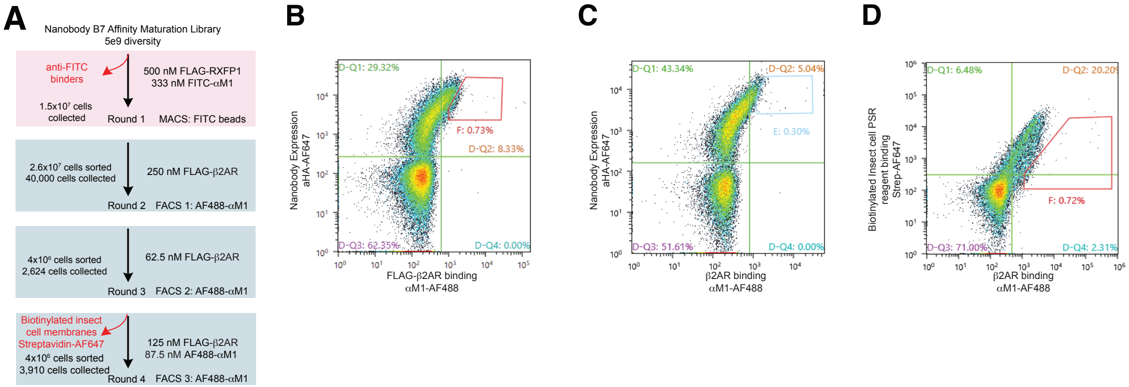


**Supp Fig 1: Affinity maturation of nanobodies that bind β2AR.** (A-D) Flowchart and Flourescence-activated cell sorting (FACS) plots for nanobody B7 affinity maturation to improve binding to β2AR.

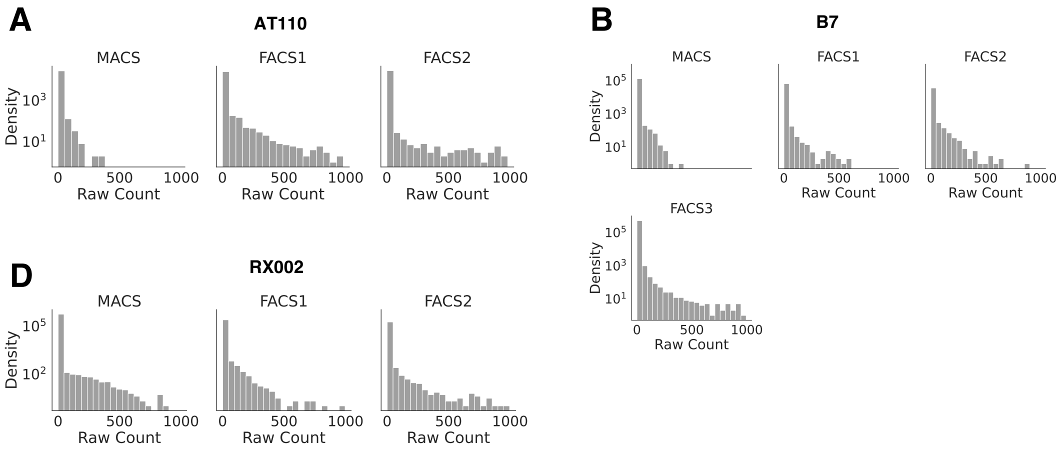


**Supp Fig 2: Count distributions of the sequencing rounds.** Histograms showing the count distributions of sequences after each sort of the affinity maturation campaigns for A) AT110, B) B7 and C) RX002. The histograms are clipped at 1000 counts to clearly show the increasing skew of the distributions as further sorts are performed.

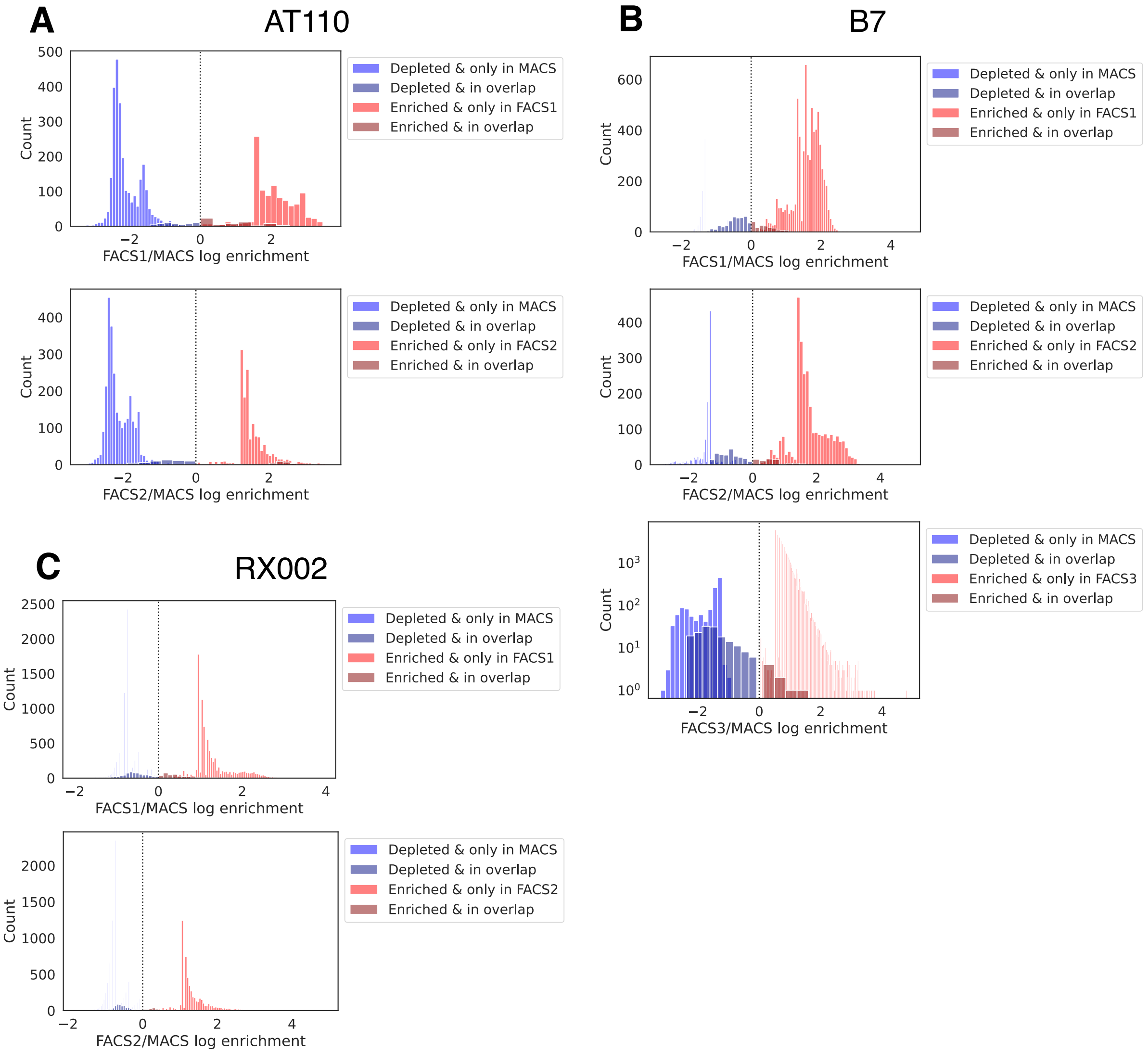


**Supp Fig 3: Enrichment distributions of the FACS rounds.** Histograms showing the distribution of log-enrichment values after each FACS sort with respect to the MACS sort of the affinity maturation campaigns for A) AT110, B) B7 and C) RX002. The distribution of sequences that are enriched and depleted after FACS are colored in red and blue respectively. Darker bars correspond to the sequences found in both rounds used to calculate the enrichment score. Lighter bars correspond to sequences only in a single round, and thus their score is determined by the sequencing count of the round they were found in and the floor abundance value used (set to 1e-6).

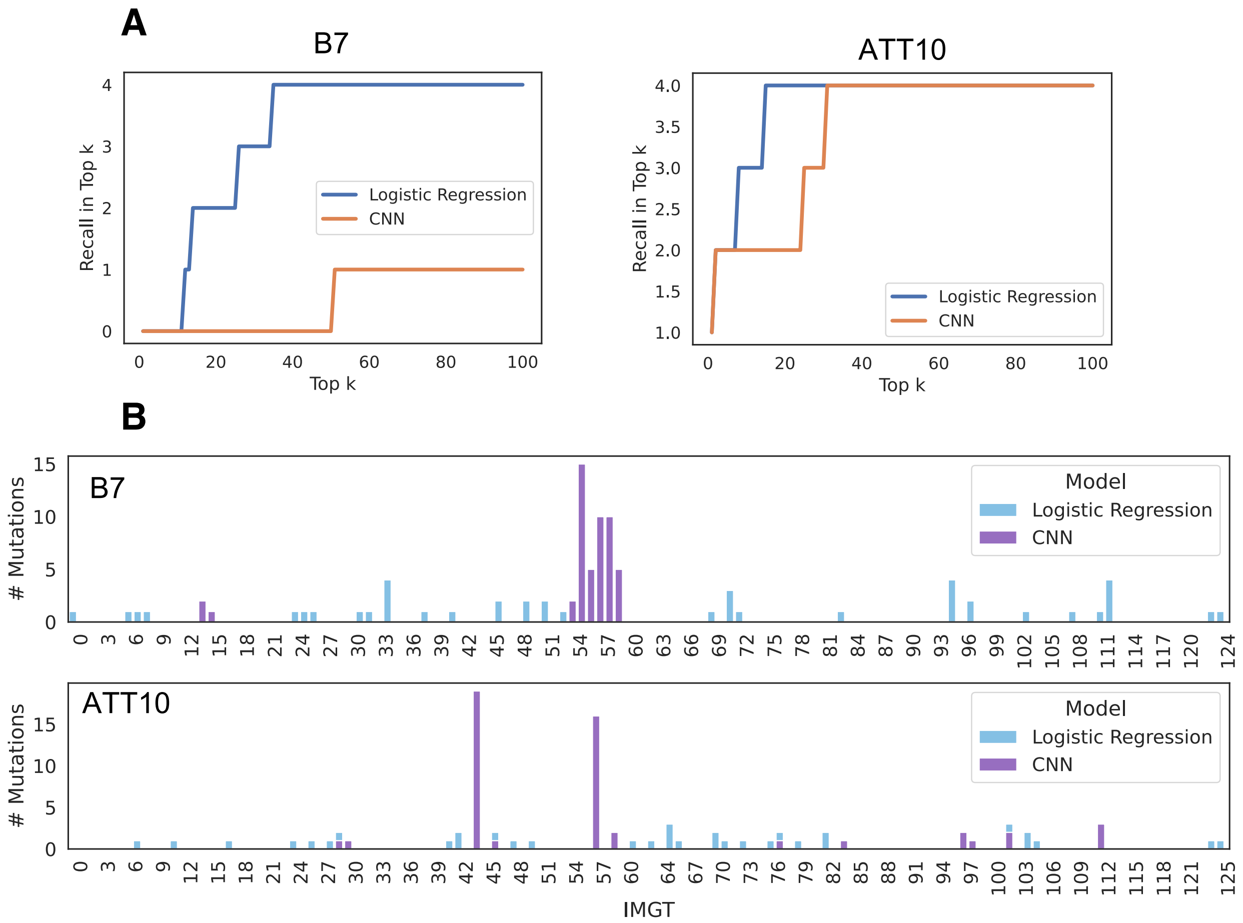


**Supp Fig 4: Comparison of top singles from a logistic regression model and CNN** A) The recall of the 4 tested mutants available for B7 and AT110 in the top k ranked single mutations from the Logistic Regression and CNN models for increasing k up to 100. B) The distribution of the IMGT sites of the top 50 mutations predicted by both models for B7 and AT110.

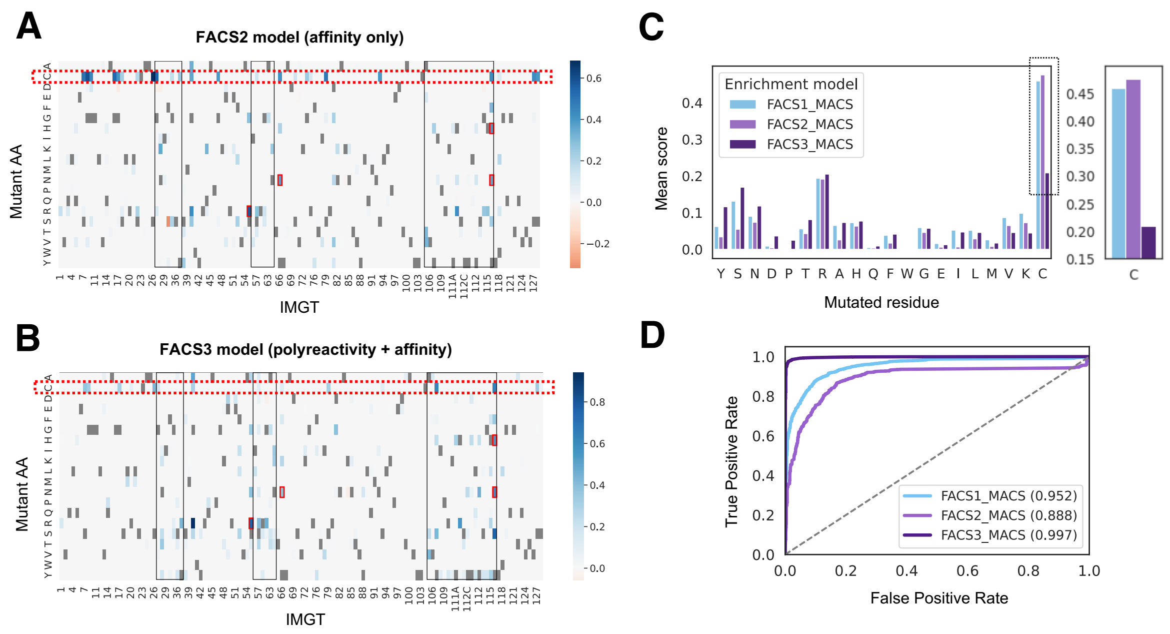


**Supp Fig 5: Analysis of polyreactivity counter-sort:** Heatmaps of scores for all single substitutions from logistic regression models trained on the A) FACS2/MACS enrichment and B) FACS3/MACAS enrichment for B7. The FACS2 sort only selected for affinity to B2AR. The FACS3 sort was a simultaneous affinity selection and polyreactivity counterselection. The scores for mutations to cysteine are highlighted with a dotted red box. Black boxes correspond to CDRs 1-3 and red boxes correspond to experimentally validated substitutions. C) (left) The average score per amino acid from the logistic regression models trained on the B7 FACS1, FACS2 and FACS3 enrichment over MACS. (right) A zoomed in view of the scores for cysteine mutations. Zoomed in portion is highlighted in black dotted lined in left panel. D) ROC-plots showing the performance of the logistic regression models trained on each round for their prediction of FACS3/MACS enrichment. The ROC-AUC for each model is shown in brackets in the legend.

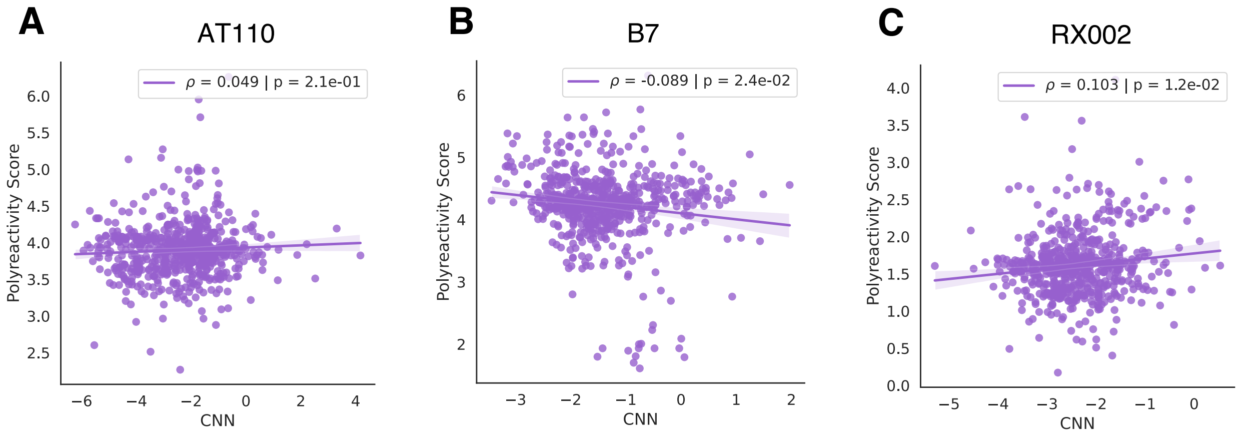


**Supp Fig 6: Affinity and Polyreactivity models have low correlation:** Scatterplots of the scores for all single substitutions from the pre-trained Polyreactivity model (Harvey et al 2022) against scores from the CNN classification models trained on the FACS1/MACS enrichment for A) AT110, B) B7, C) RX002 and D) LysoNbV1. LpoB was not analyzed in this way as it is not a nanobody. Each point corresponds to a single substitution of the corresponding binder. A line of best fit is plotted with bootstrap estimated 95% confidence intervals as shaded intervals. The spearman correlation and pvalue for the correlation is shown in scatter legend.

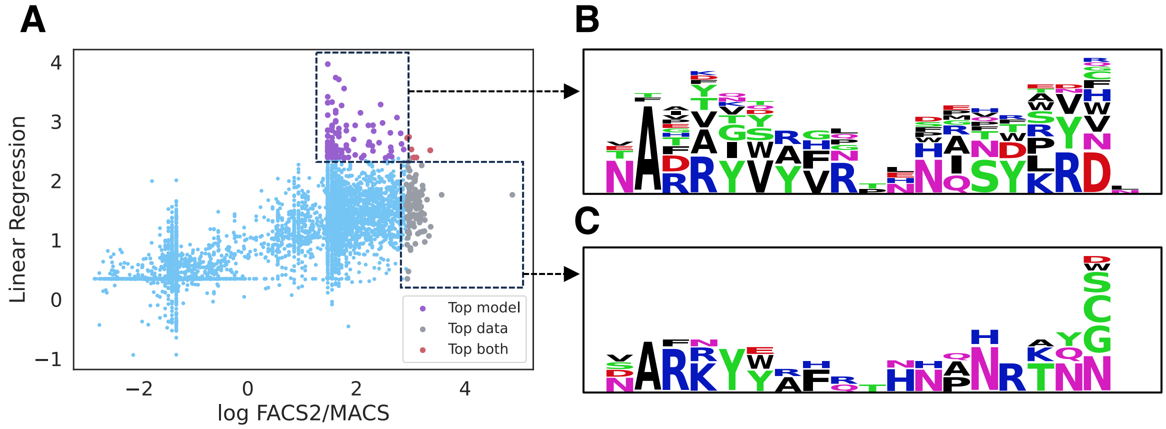


**Supp Fig 7: Affinity models suggest different top sequences from the experiment:** A) Scatterplot of scores from a linear regression model compared to the continuous FACS2/MACS enrichment score for sequences in the B7 campaign. The top 100 sequences by model score and FASC2 enrichment are colored in purple and gray respectively. Sequences in both sets are colored red. B-C) Sequence logos of the CDR3 region for the top 100 sequences under the model score (B) and enrichment from the experiment (C). Logos are plotted by calculating the amino acid frequencies for each site in the CDR3 (excluding gaps). The amino acid corresponding to the WT residue at that site is also omitted from the logo. Thus, the height of each residue corresponds to the relative probability of a substitution to that amino acid, and the height of the whole column corresponds to the probability of any substitution at that site.

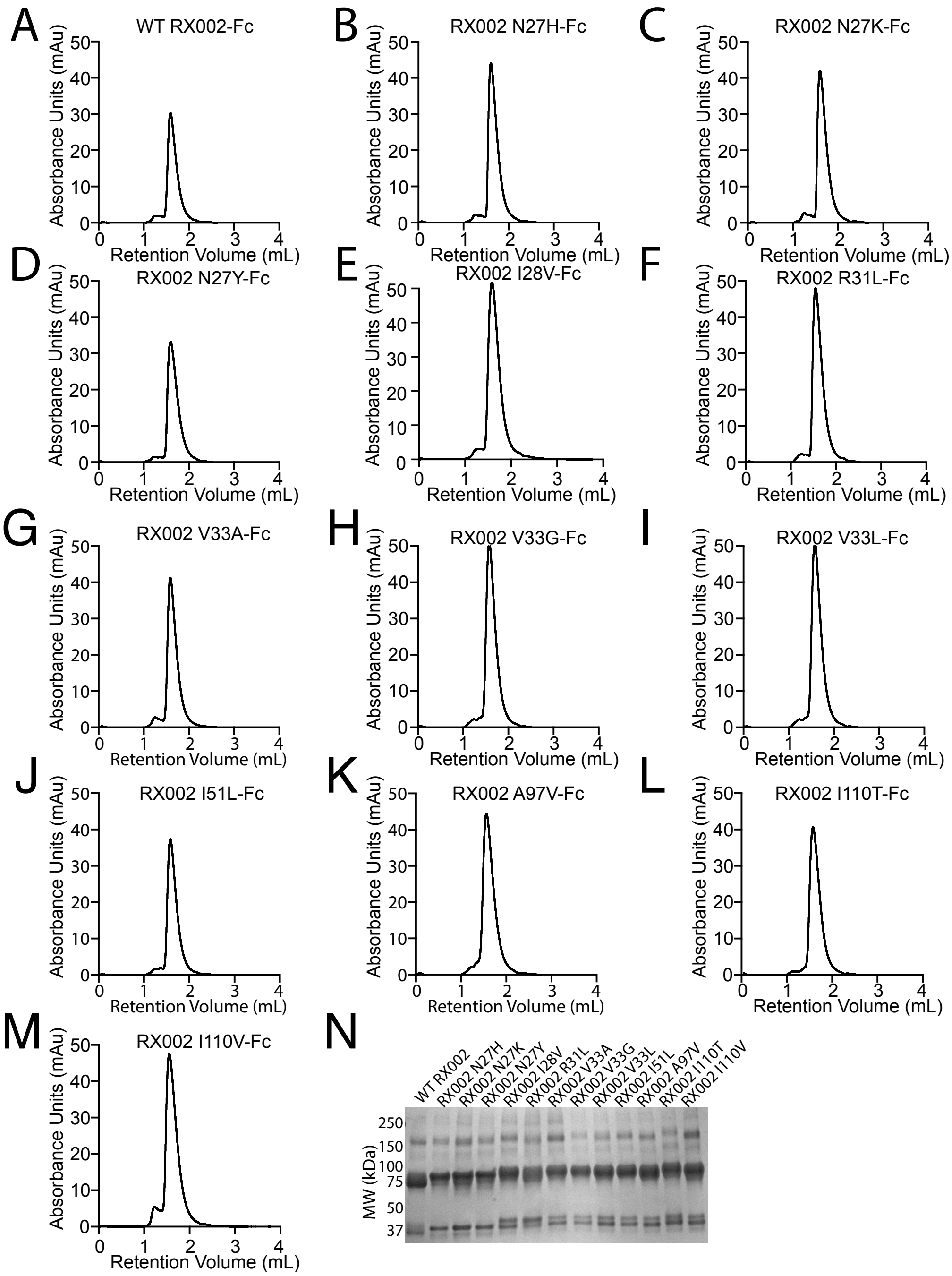


**Supp Fig 8: Size exclusion chromatograms for RX002 mutants:** A-M) The quality of purified, isogenic FC-fusion versions of selected RX002 mutants were measured by size exclusion chromatography (SEC). Each protein had only one peak demonstrating pure product. N) Western blot showing expression of RX002 at a MW of 37 kDa. One band is observed for the N27 mutants with two bands observed for WT and the other mutants. We hypothesize this is due to a disruption of a glycosylation site by mutating residue N27.

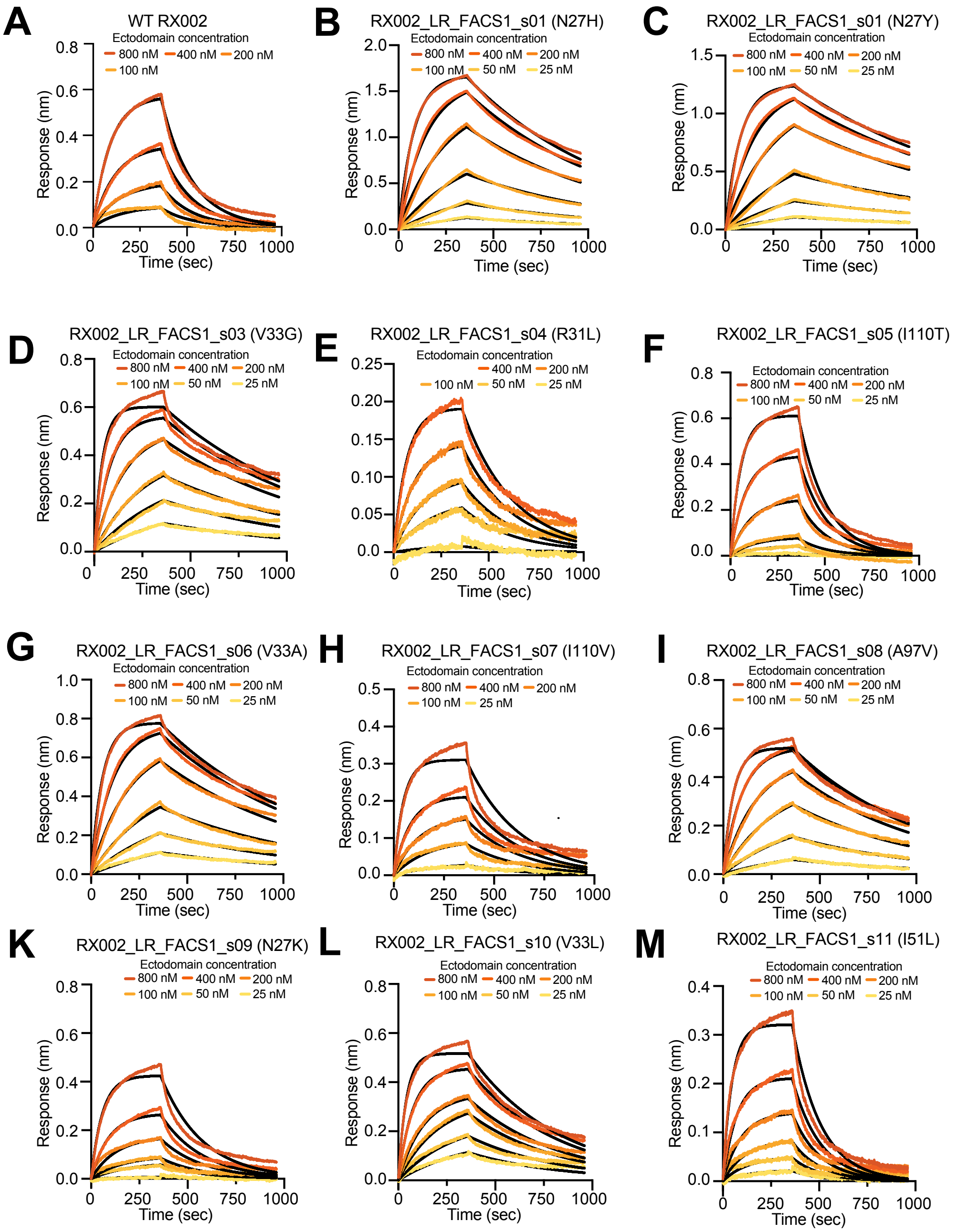


**Supp Fig 9: Dissociation curves for RX002 mutants measure by BLI:** A-F) Dissociation curves for multiple concentrations of each selected RX002 mutant binding to purified RXFP1 target protein. Black lines correspond to parametric fit for each with the orange gradient corresponding to binder concentration.

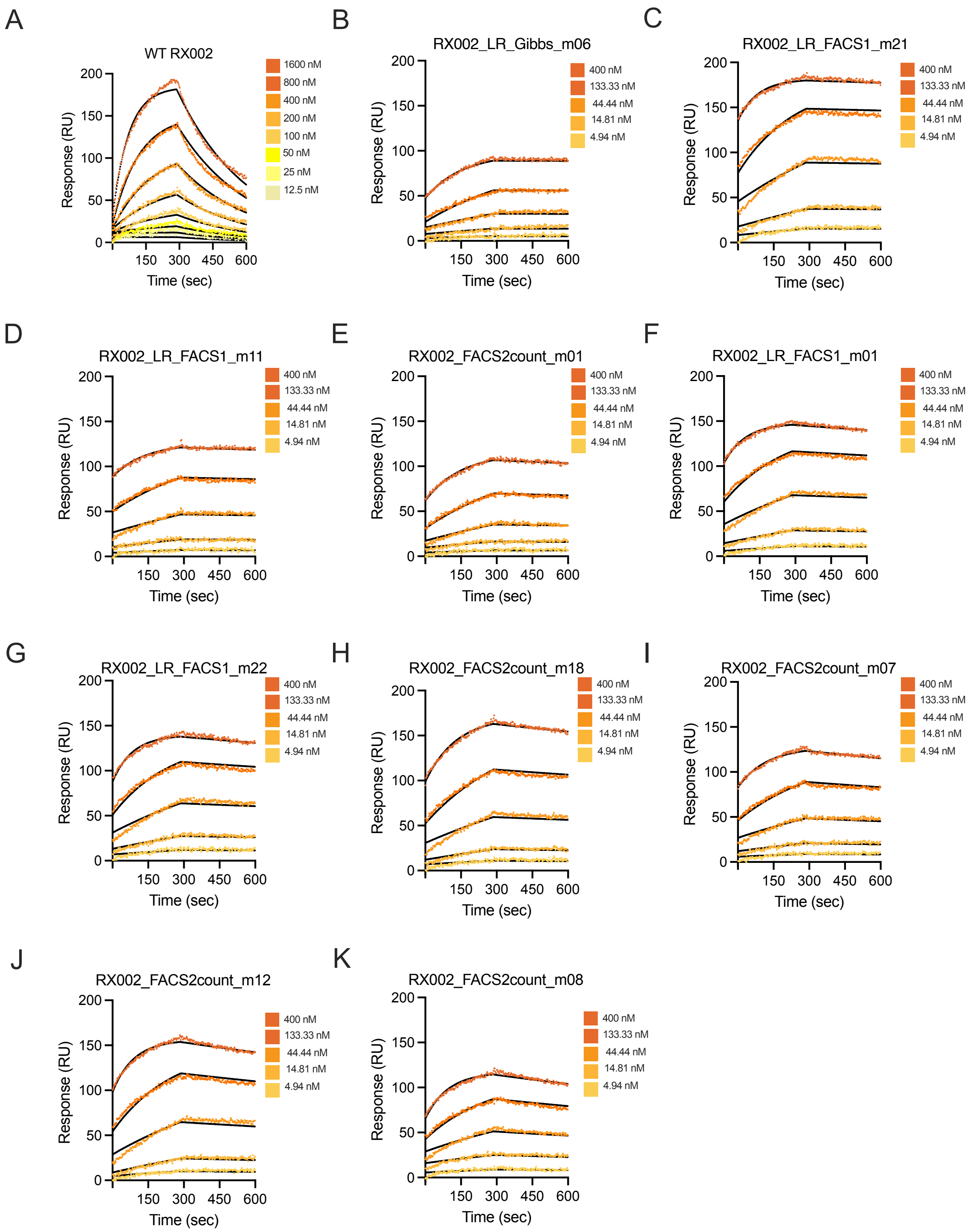


**Supp Fig 10: Dissociation curves for top designed RX002 multiple mutants measured by SPR:** A-K) Dissociation curves for binding to purified RXFP1 target protein of multiple concentrations of WT RX002 and the 10 tested multiple mutants with lowest $K_D$. Black lines correspond to parametric fit for each with the orange gradient corresponding to binder concentration.

| Nanobody | Round | # Total Reads | # Unique Sequences | # Sequences above Threshold |
|---|---|---|---|---|
| *AT110* | MACS | 118,429 | 26,516 | 2,805 |
| | FACS1 | 148,783 | 24,648 | 1,172 |
| | FACS2 | 278,191 | 27,053 | 1,539 |
| *B7* | MACS | 241,225 | 134,340 | 1,433 |
| | FACS1 | 230,676 | 66,641 | 7,160 |
| | FACS2 | 168,987 | 37,344 | 3,131 |
| | FACS3 | 1,521,316 | 520,440 | 42,012 |
| *RX002* | MACS | 922,141 | 555,762 | 8,543 |
| | FACS1 | 554,583 | 246,068 | 9,331 |
| | FACS2 | 430,340 | 180,670 | 6,126 |

**Supp Table 1.** Sequencing dataset sizes. Summary count statistics of the sequencing from the affinity maturation campaigns used in this study. We use a threshold of 5 counts for retaining sequences for training machine learning models.

| Nanobody | $R_0/R_1$ | $R_0 \cap R_1$ | % | $R_0^* \cap R_1^*$ | % | $R_0^* \cup R_1^*$ | (#depleted, #enriched) |
|---|---|---|---|---|---|---|---|
| *AT110* | FACS1/MACS | 1173 | 2.35% | 111 | 2.87% | 3866 | (2747, 1119) |
| | FACS2/MACS | 1166 | 2.23% | 83 | 1.95% | 4261 | (2783, 1478) |
| *B7* | FACS1/MACS | 5853 | 3.00% | 561 | 6.98% | 8032 | (1280, 6752) |
| | FACS2/MACS | 2902 | 1.72% | 284 | 6.64% | 4280 | (1365, 2915) |
| | FACS3/MACS | 1656 | 0.25% | 170 | 0.39% | 43275 | (1516, 41759) |
| *RX002* | FACS1/MACS | 12922 | 1.64% | 1016 | 6.03% | 16858 | (8154, 8704) |
| | FACS2/MACS | 12911 | 1.78% | 798 | 5.75% | 13871 | (8202, 5669) |

**Supp Table 2: Overlap statistics and train set sizes.** $R_0$ and $R_1$ describe which two rounds we are considering the overlap between. $R_0 \cap R_1$ is the number of overlapping unique sequences found in both rounds with the % reflecting what this overlap is as a proportion of the union of the sets. $R_0^*$ and $R_1^*$ describe the sequencing sets after filtering by removing low count sequences. $R_0^* \cup R_1^*$ is the total number of sequences that pass the filtering threshold in either of the rounds. This is the training data size for that round. The number of sequences of each label in each training set is shown in the last column.

| Count Threshold | ROC-AUC | NDCG@100 |
|---|---|---|
| 1 | 0.9254 | 0.812 |
| 2 | 0.9418 | 0.855 |
| 3 | 0.9467 | 0.852 |
| 4 | 0.9507 | 0.848 |
| 5 | 0.9519 | 0.857 |

**Supp Table 3: Effect of count threshold on model performance.** The ROC-AUC of FACS2/MACS enrichment prediction and the NDCG@100 for ranking known optimizing mutations for varying values of the count threshold used to filter out low count sequences. Results here are for Logistic Regression models trained on the FACS1/MACS enrichment from the AT110 campaign. Higher count thresholds increase the performance of the model, likely due to increasing data quality of the training data.

| Nanobody | Kd (M) Rep1 | Kd (M) Rep2 | Kd (M) Rep3 | Kd (M) Mean | Fold Change over WT |
|---|---|---|---|---|---|
| WT RX002 | 2.46E-06 | 5.73E-07 | 1.04E-06 | 1.3596E-06 | 1.00 |
| RX002 N27H | 9.25E-08 | 8.6E-08 | 9.41E-08 | 9.0867E-08 | *14.96 |
| RX002 N27Y | 6.15E-08 | 5.5E-08 | 4.82E-08 | 5.4887E-08 | *24.77 |
| RX002 V33G | 4.36E-08 | 2.9E-08 | 4.11E-08 | 3.7887E-08 | *35.89 |
| RX002 R31L | 5.48E-08 | 1.38E-07 | 1.19E-07 | 1.0376E-07 | *13.10 |
| RX002 I110T | 4.79E-07 | 1.68E-07 | 2.9E-06 | 1.1825E-06 | 1.15 |
| RX002 V33A | 6.48E-08 | 5.07E-08 | 4.88E-08 | 5.476E-08 | *24.83 |
| RX002 I110V | 2.2E-07 | 1.76E-07 | 5.37E-07 | 3.111E-07 | 4.37 |
| RX002 A97V | 4.19E-08 | 6.77E-08 | 5.81E-08 | 5.59E-08 | *24.32 |
| RX002 N27K | 1.49E-07 | 2.95E-07 | 3.41E-07 | 2.6157E-07 | 5.20 |
| RX002 V33L | 1.72E-07 | 7.8E-08 | 1.09E-07 | 1.1938E-07 | *11.39 |
| RX002 I51L | 3.68E-07 | 3.82E-07 | 5.4E-06 | 2.0504E-06 | 0.66 |

**Supp Table 4: Binding affinities for RX002 model selected substitutions.** The $K_d$ (M) for the 10 model selected single substitutions of nanobody RX002 was measured by biolayer inferometry (BLI), along with the WT RX002 as positive control and a known deleterious mutation (I51L) as a negative control. The fold change of the mutant affinity over the WT is shown in the last column. Mutants with at least a 10-fold affinity increase are starred(*).

| General Primers | | |
|---|---|---|
| epPCR_MFAlpha_Fwd | GTTCAATTGGACAAGAGAGAAGCT | Integrated DNA Technologies |
| epPCR_HA_Rvs | GAACATCGTATGGGTAGGATCC | Integrated DNA Technologies |
| epPCR_Lib_fwd | ATCGCTGCTAAGGAAGAAGGTGTTCAATTGGACAAGAGAGAAGCT | Integrated DNA Technologies |
| epPCR_Lib_rvs | GGGTGAGGATGTTTGAGCGTAATCTGGAACATCGTATGGGTAGGATCC | Integrated DNA Technologies |
| epPCR_Gib_1 | ACCTTCTTCCTTAGCAGCGAT | Integrated DNA Technologies |
| epPCR_Gib_2 | TACGCTCAAACATCCTCACCC | Integrated DNA Technologies |
| pGAL nanobody sequencing | AATATACCTCTATACTTTAACGTC | Integrated DNA Technologies |

| Nanobody NGS Primers | | |
|---|---|---|
| Primer Set | Forward | Reverse |
| NGS #1 | **TCGAAG**GTTCAATTGGACAAGAGAGAAGCT | CTTCGAGTAATCTGGAACATCGTATGGGTA |
| NGS #2 | **ACCTGA**GTTCAATTGGACAAGAGAGAAGCT | TCAGGTGTAATCTGGAACATCGTATGGGTA |
| NGS #3 | **GCAATC**GTTCAATTGGACAAGAGAGAAGCT | GATTGCGTAATCTGGAACATCGTATGGGTA |
| NGS #4 | **GAGATT**GTTCAATTGGACAAGAGAGAAGCT | TAATCTGTAATCTGGAACATCGTATGGGTA |
| NGS #5 | **ATCACG**GTTCAATTGGACAAGAGAGAAGCT | CGATGTGTAATCTGGAACATCGTATGGGTA |
| NGS #6 | **TTAGGC**GTTCAATTGGACAAGAGAGAAGCT | TGACCAGTAATCTGGAACATCGTATGGGTA |
| NGS #7 | **ACAGTG**GTTCAATTGGACAAGAGAGAAGCT | GCCAATGTAATCTGGAACATCGTATGGGTA |
| NGS #8 | **CAGATC**GTTCAATTGGACAAGAGAGAAGCT | ACTTGAGTAATCTGGAACATCGTATGGGTA |
| NGS #9 | **ATGCAC**GTTCAATTGGACAAGAGAGAAGCT | TACATGGTAATCTGGAACATCGTATGGGTA |

Error Prone PCR Library Generation Protocol

**Materials needed:**

Agilent Technologies GeneMorph II Random Mutagenesis Kit (Agilent Cat# 200550)
KOD Xtreme Polymerase (MilliporeSigma Cat# 71-975-3)
QIAquick PCR Purification Kit (Qiagen Cat# 28104)
QIAquick Gel Extraction Kit (Qiagen Cat# 28704)
NEBuilder HiFi DNA Assembly Master Mix (NEB Cat# 2621L)
NheI HF restriction digest enzyme (NEB Cat# R3131S)
BamHI HF restriction digest enzyme (NEB Cat# R3136S)
PureLink HiPure Plasmid Filter Midiprep Kit (Invitrogen, Cat# K210014)
PureLink HiPure Precipitator Module (Invitrogen, Cat# K2100-22)
BJ5465 Yeast (from ATCC)
epPCR_MFAlpha_Fwd Primer: GTTCAATTGGACAAGAGAGAAGCT
epPCR_HA_Rvs Primer: GAACATCGTATGGGTAGGATCC
epPCR_Lib_fwd Primer: ATCGCTGCTAAGGAAGAAGGTGTTCAATTGGACAAGAGAGAAGCT
epPCR_Lib_rvs Primer: GGGTGAGGATGTTTGAGCGTAATCTGGAACATCGTATGGGTAGGATCC
epPCR_Gib_1 Primer: ACCTTCTTCCTTAGCAGCGAT
epPCR_Gib_2 Primer: TACGCTCAAACATCCTCACCC
pGAL Sequencing Primer: TACGCTCAAACATCCTCACCCC
100% ethanol pre-chilled to -20 °C
70% ethanol pre-chilled to 4 °C
3 M sodium acetate pH = 5.2

**Protocol:**

**Error Prone PCR:**

Using the epPCR_MFAlpha_Fwd and epPCR_HA_Rvs primers and the nanobody plasmid that you want to mutagenize, perform error prone PCRs according to the GeneMorph II kit instructions. In particular, choose 4-5 different starting concentrations of plasmid and cycle numbers to achieve different plasmid error rates. The PCR conditions needed are as follows:

| | | |
|---|---|---|
| 42.5 µL Nanobody DNA + dI $H_2O$ | 1). 95°C | 2 minutes |
| 5 µL 10x Mutazyme II Rxn Buffer | 2). 95°C | 30 seconds |
| 1 µL 40 mM dNTPs (for 200 µM final) | 3). 50°C | 30 seconds |
| 0.5 µL primer mix (250 ng/µL of each primer) | 4). 72°C | 1 minute |
| 1 µL Mutazyme DNA polymerase | Repeat steps 2-4 30x times | |
| | 5). 72°C | 10 minutes |

To generate a library of sufficient diversity, you want to have pools of mutagenized plasmid with different error rates (i.e. 1 mutation, 2 mutations, 4 mutations, 6 mutations, etc). The lower the plasmid concentration you use and higher the PCR cycle count, the greater the error rate. These are the error rates for the nanobody B7 β2AR library using the listed concentration of plasmid and cycle number. Note they are lower than what the kit says you should get. Also, this refers to total plasmid DNA not target DNA.

40 ng with 30 cycles: 2.16 +/- 0.98
15 ng with 30 cycles: 4.72 +/- 2.32
5 ng with 30 cycles: 5 +/- 1.25
2 ng with 35 cycles: 6.54 +/- 3.56

After the EP-PCR, clean up each sample using the PCR purification kit. Always include the optional 10 µL sodium acetate pH = 4 and elute with 35 µL dI $H_2O$.

**Gibson Cloning to Check Error Rate**
Now that you have mutated inserts, check the error rates before building the library. Amplify the PYDS2.0 vector (length 9,921 base pairs) using primers epPCR_Gib_1_55 and epPCR_Gib_2 Primer using the KOD xtreme polymerase and the following PCR conditions per reaction:

6 µL Sterile H2O
12.5 µL 2x xtreme buffer
5 µL 2 mM dNTPs
0.3 µL forward primer
0.3 µL reverse primer
0.6 µL KOD xtreme Polymerase
0.4 µL 62.5 ng/µL PYDS2.0 plasmid

1). 94°C 2 minutes
2). 98°C 10 seconds
3). 55°C 30 seconds
4). 68°C 9 minute 20 seconds
repeat steps 2-4 25x times
5). 68°C 10 minutes

After the PCR, check that the product is the right size by running an agarose gel (0.8 mg agarose in 50 mL TAE). If it is the right size, add 1 µL DPN1 and incubate overnight at 37°C.

Use the Gibson Template and the HiFi DNA Assembly Mastermix to check the error rate. The reaction volume should be 10 µL total (5 µL of the Assembly Mastermix and 5 µL DNA/$H_2O$). Take 2 µL and transform XL1-Blue competent cells. Incubate DNA with cells for 30 minutes and then recover at 37°C using 250 µL SOC media for >2 hours. The following morning, submit the plates to Genewiz for combined miniprep and sequencing ( 10-15 colonies is standard to submit for sequencing). Submit 5 µL of 5 mM pGAL sequencing primer for every colony you'd like sequenced. The pGAL sequencing primer will sequence nanobody inserts in both pYDS/pYDS2.0 plasmids.

**PCR Amplification**
For DNA pools that have acceptable mutation rates, a second PCR step is needed to add homology arms for yeast electroporations (homology arms must be >30 bp for efficient yeast homologous recombination). Additionally, since transformation of yeast by electroporation requires a large amount of insert and restriction digested plasmid, we need to amplify the error prone PCR product using Q5 master mix and primers epPCR_Lib_fwd and epPCR_Lib_rvs. Here are conditions you need per reaction (set up a considerable number of reactions to have enough DNA for a large library).

25 µL Q5 Master Mix

1). 98°C 30 seconds

| | |
|---|---|
| ~600 nM epPCR_Lib_fwd Primer | 2). 98°C 10 seconds |
| ~600 nM epPCR_Lib_rvs Primer | 3). 72°C 30 seconds |
| 20 ng DNA | 4). 72°C 1 minute |
| ddH20 to 50 μL | repeat steps 2-4 30x times |
| | 5). 72°C 2 minutes |

Following the PCR amplification, perform gel electrophoresis on the PCR product. Make a agarose gel containing 0.8 g agarose in 50 mL TAE buffer with 5 μL Cybersafe stain. Gel running conditions are 100V for 25 minutes. Cut out the nanobody bands of the appropriate size (474 base pairs) and use the gel extraction kit to isolate the DNA, using ice cold isopropanol and letting the PE buffer sit for 5 minutes prior to removal to reduce salt (which will cause arcing during the transformation). Note that you may also wish to perform a PCR purification as opposed to a gel extraction at this step because your yield will be higher. Once you have all of your inserts amplified, you should perform an ethanol precipitation to ensure the electroporator does not arc (see below).

**Digesting the PYDS2.0 Plasmid**

To build a large library using yeast electroporation, you need to digest significant amount of pYDS2.0 vector (which contains a TEF promoter and nourseothricin resistance cassette as a selectable marker). First, midiprep or maxiprep the PYDS2.0 vector, and use a precipitator kit or ethanol precipitation to make sure there is no salt left. Then, digest at least 100 μg of pYDS2.0 using the following reaction scheme.

100 μg pYDS2.0
40 μL cutsmart
20 μL NheI-HF
20 μL BamHI-HF
Remainder DI $H_2O$ for 400 μL total

Incubate at 37°C for an hour and a half. Then do the PCR purification kit (~5 columns for 100 μg). Run an agarose gel to make sure the cut and uncut PYDS2.0 different in size and the cut PYDS2.0 is the appropriate size.

For larger quantities of DNA (> 1 μg), the DNA will substantially add to the ionic strength of the solution – you must make sure the DNA is salt free by ethanol precipitation:

**Ethanol Precipitation Protocol**

1. Check DNA concentration by nanodrop to be able to calculate recovery at the end.
2. To tube containing x μL DNA, add 0.1x volume of 3 M sodium acetate pH 5.2. It is helpful if DNA is less than 400 μL so you can do everything in one tube.

3. Add 2.5x volumes ice-cold 100% ethanol (should be pre-chilled at -20 °C).
4. Mix with pipette and incubate in -20 °C freezer for 1 hour up to overnight (longer tends to be better for dilute and small pieces of DNA).
5. Centrifuge at maximum speed (≥18,000 *x g*) for 30 minutes at 4 °C. It is helpful to note the tube's orientation in the centrifuge–e.g., by having the hinge face out—so you can locate the pellet later.
6. Carefully remove supernatant with pipette. Pellet should be small and clear.
7. Add 200 µL of cold 70% ethanol (should be pre-chilled at 4 °C). Pipette up and down until pellet dislodges from the side of the tube. Centrifuge at max speed for 10 minutes at 4 °C.
8. Repeat wash step (steps 6 and 7) one more time.
9. After second wash, remove as much supernatant as possible. Open tube and allow pellet to air dry (this works at room temperature but can also be done at 37 °C).
10. Resuspend pellet in desired volume of water/buffer and mix by pipetting.
11. Measure concentration by nanodrop and calculate recovery (should be ≥90%).

**Transforming Yeast by Electroporation**
Day 1: start 5 mL starter culture in YPAD and grow overnight
Day 2: inoculate 100 mL starter culture in YPAD
When yeast reach OD of 1.5, transfer 50 mLs to falcons and pellet 2000xg for 5 min, discard supernatant. Each electroporation cuvette will give you approximately 2e7 transformations so you can back calculate how many yeast you will need for a certain library size.
Resuspend each falcon tube with 25 mL sterile 100 mM Lithium Acetate. Shake vigorously.
Add 0.25 mL sterile-filtered fresh 1 M DTT to each tube.
Loosen cap and attach with tape, shake cells at 30 °C for 10 min in incubator.

Perform all additional steps on ice
Tighten caps, pellet at 2000xg for 5 min. Discard supernatant.
Resuspend each falcon with 25 mL ice-cold sterile water and shake vigorously.
Pellet 2000xg 5 min, discard supernatant
Resuspend cell pellet in vector/insert mix- for 50 rxns use 50 µg cut vector, 250 µg insert- 2.5 mL total

Place 50 µL in square wave electroporator- 500V, 15 ms pulse, plate
Estimate the diversity of the yeast post transformation by calculating how many cells you electroporated, and then plating sequential dilutions on YPAD plates.

After electroporating, recover cells in YPD without shaking at 30°C for 45-60 minutes. Plate out different dilutions of yeast to calculate the diversity of the library. It is very important to do this before adding the selection antibiotic if you are using pYDS2.0. If you are using an auxotrophic selection system, do the colony dilutions after washing into dropout media. Add selection antibiotic (100 ug/mL Nourseothricin) or wash into dropout medium, as appropriate. Recover for 2 days at 30°C with shaking at 180xRPM at 30°C. Count colonies on dilution plates after 2 days recovery without shaking at 30°C.

### Freezing Down Post Electroporation

1). Yglc4.5 –Trp (1 liter): For amplifying library

i. 3.8 g -Trp drop-out media supplement (US Biological)
ii. 6.7 g Yeast Nitrogen Base
iii. 10.4 g Sodium Citrate
iv. 4.4 g Citric Acid Monohydrate
v. 10 mL Pen/Strep (10,000 units/mL stock)
vi. 20 g glucose

1. Adjust pH to 4.5, sterifilter (alternatively prepare media without glucose or pen/strep, autoclave and add sterile 20% glucose solution an antibiotic after autoclaving)

2). Measure the OD of your yeast and do the conversion with $1.5x10^7$ conversion factor.

3). Spin down 10x diversity of library at 3500 rpm for 4 minutes in Eppendorf tubes. Save A LOT of them because you'll never know when you'll want/need to go back to them.

4). Make Yglc4.5-Trp + 10% DMSO. Make the Yglc4.5-Trp first and then take 9 mL and add 1 mL fresh DMSO to it.

5). Remove the supernatant from the Eppendorf tubes so that only the yeast pellets are left. Resuspend in 750 μL of Yglc4.5-Trp + 10% DMSO.

6). Put all the tubes in a Mr. Freezey chamber and put it in the -80°C. Wait a day and then store them long term
